# Molecular roadmap of the healthy stress response in the mouse hippocampus

**DOI:** 10.1101/2021.03.26.436970

**Authors:** Lukas M. von Ziegler, Amalia Floriou-Servou, Rebecca Waag, Rebecca R. Das Gupta, Oliver Sturman, Katharina Gapp, Christina A. Maat, Tobias Kockmann, Han-Yu Lin, Sian N. Duss, Mattia Privitera, Laura Hinte, Ferdinand von Meyenn, Hanns U. Zeilhofer, Pierre-Luc Germain, Johannes Bohacek

**Author notes:** **Corresponding Author:** Johannes Bohacek.

## Abstract

The acute stress response mobilizes energy to meet situational demands and re-establish homeostasis. However, little is known about the nature and dynamics of the underlying molecular cascades. We used a brief forced swim exposure to trigger a strong stress response in mice, which transiently increases anxiety, but does not lead to lasting maladaptive behavioral changes. Using multiomic profiling we characterize the stress-induced molecular events in the hippocampus over time, across molecular scales, and down to the level of cell types and single cells. Our results unveil the complexity and specificity of the healthy stress response, with widespread changes in protein phosphorylation and gene transcription, but tightly regulated protein translation. All observed molecular events resolve efficiently within 4 hours after initiation of stress. The data are accessible through an interactive web portal, creating an extensive molecular resource that will help understand how stress impacts brain function in health and disease.

## Introduction

The acute stress (AS) response enhances the chance of survival by mobilizing organismal energy resources to meet situational demands (Cannon, 1915; Floriou-Servou et al., 2021; Ulrich-Lai and Herman, 2009). When the stressor has subsided, successful stress management requires the efficient termination of the stress response to avoid stress-related wear and tear (De Kloet et al., 2005; McEwen, 1998; Sapolsky, 1996). Notably, the majority of individuals can successfully cope with stressful challenges and maintain mental health even in the face of severe stressors (Boden and McLeod, 2015; Bonanno et al., 2011; Kalisch et al., 2017). However, when AS becomes too intense, or occurs repeatedly (i.e. chronic stress), it can overwhelm the “healthy stress response” and give rise to neuropsychiatric diseases such as post-traumatic stress disorder, anxiety and depression (McEwen and Akil, 2020; Musazzi et al., 2017a; Tost et al., 2015). To understand the underlying mechanisms, many studies have assessed how chronic stressors impact the genome-wide molecular landscape in the brain, particularly in the hippocampus (HC) (Bagot et al., 2016; Cerniauskas et al., 2019; Gray et al., 2014; Hamilton et al., 2020; Lepack et al., 2016; Misiewicz et al., 2019; Peña et al., 2019; Sun et al., 2015; Wei et al., 2020). However, much less is known about the molecular changes triggered by AS, and how they unfold dynamically over time and across multiple molecular levels. The few available studies have only catalogued gene expression changes at single time points after AS exposure (Floriou-Servou et al., 2018; Gray et al., 2014; Roszkowski et al., 2016; Stankiewicz et al., 2015; Terenina et al., 2019; Tsolakidou et al., 2008). Thus, we only have a fragmented picture of how AS affects the molecular machinery in the HC, and many key questions remain unanswered. First, how do stress-induced effects on transcription evolve over time and how long do they persist? Second, in which cell types do these changes occur? Third, which molecular changes occur upstream of transcription, at the level of protein phosphorylation and transcription factor activity? Fourth, which molecular changes occur downstream of transcription, at the level of translation and protein regulation? Here, we use a multiomic approach to dissect the healthy stress response across molecular scales, by cataloguing the stress-induced changes at the level of the phospho-proteome, transcriptome, translatome and proteome. We look across cell types and profile the molecular changes over time, to reveal how an organism mounts a peak stress response and then reestablishes homeostasis after a brief, intensely stressful experience.

## Results

### Stress-induced effects on behavior

To induce AS, we chose the forced swim stress model, which strongly activates the sympathetic nervous system and the HPA axis (Floriou-Servou et al., 2018; Gray et al., 2014; Mifsud and Reul, 2016; Roszkowski et al., 2016). Here we first confirmed previous work showing that mice are anxious and less exploratory shortly after an AS exposure (Korte and De Boer, 2003; Sturman et al., 2018). When tested 45 min after an AS in the open field test (OFT), mice travelled less distance, spent less time in the center, and performed fewer supported and unsupported rears (Figure S1B). However, when tested 2, 4 or 24 hours after the initiation of stress, these changes largely disappeared and stress-exposed mice appeared behaviorally similar to unstressed control mice in the OFT (Figure S1C) and indistinguishable to controls in the elevated plus maze (EPM) (Figure S1D). To further test whether behavior indeed normalized completely after AS exposure, we used a deep learning approach for high-precision tracking of mice (Mathis et al., 2018), as previously described (Sturman et al., 2020), in combination with unsupervised k-means clustering. Such approaches are exquisitely sensitive to detecting even subtle behavioral changes in mice (Hsu and Yttri, 2019; Wiltschko et al., 2020). Using this novel approach, we repeated the behavior experiment and tested mice 45 min after AS on the OFT, and 24 hours later on the OFT and EPM (Figure 1A). We identified clearly distinguishable clusters of behavior cross-validated with t-SNE (Figure 1B) and visual inspection (see Supplementary Videos S4). It confirmed widespread behavioral changes shortly after stress exposure (Figure 1D top), but detected no major changes in behavior the next day, neither on the OFT (Figure 1D bottom) nor on the EPM (Figure 1E). A correlation analysis revealed that independent clusters were identified, but also some that correlated to each other (Figure 1C), in line with factor analyses of OFT data, showing that different behaviors are related to each other (Crusio, 2001; Sturman et al., 2018; Whimbey and Denenberg, 1967). Closer visual inspection of altered clusters revealed that clusters 9, 10 and 14 contain many supported rears, clusters 1 and 11 contain walking behaviors in the center, and cluster 18 contained grooming behavior. Classical readouts from the same experiment indicated reduced distance and supported rears after 45 min in stressed mice in the OFT, but no group differences after 24 hours in both the OFT and EPM (Figures S1E-G). The swift and almost complete recovery within 24 hours indicates that - despite a strong initial response - mice cope successfully with this potentially life-threatening experience, and show no long-lasting alterations in anxiety-related behavior. Therefore, the acute swim stress model appears to be suitable for studying successful, “healthy” stress coping mechanisms and the underlying molecular changes in the brain.

**Figure 1:**
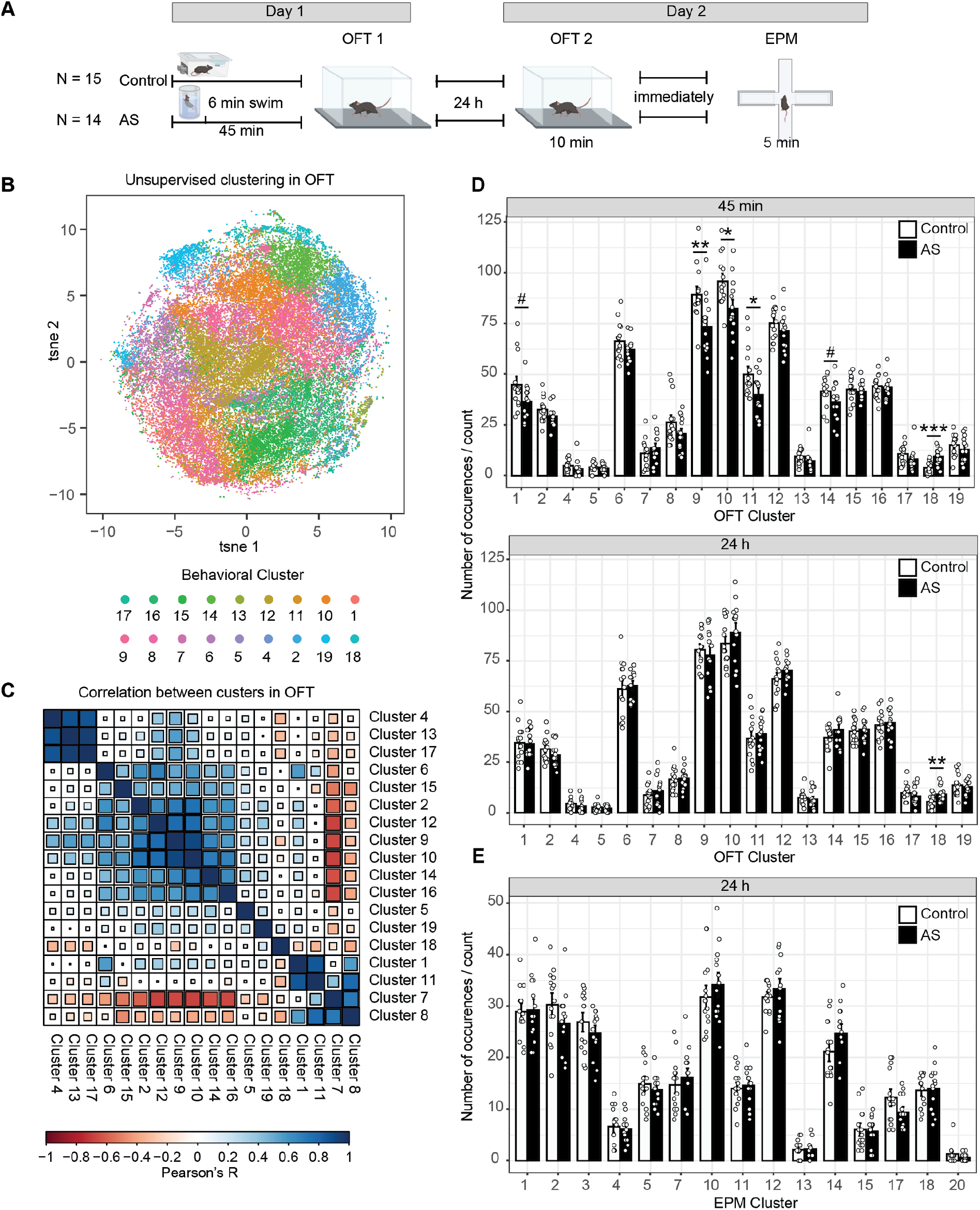
Acute stress (AS) exerts immediate but transient behavioral effects. **A)** Experimental overview to assess behavioral changes 45 min and 24 hours after AS **B)** behavioral clustering cross validated with t-sne demonstrates a reasonable separation of k-means clusters in OFT. Behavioral sequences are plotted by tsne coordinates and colored by cluster numbers independently obtained with k-means. **C)** Correlation between clusters in the OFT. **D)** OFT clusters altered by AS in the OFT: *top:* 45 min after swim, many clusters show significant differences between control and stressed mice. *bottom:* 24 hours after swim, only a single cluster shows significant differences between control and stressed mice. **E)** EPM clusters show no difference between controls and 24 hours post-swim. #p < 0.1, *p < 0.05, **p < 0.01, ***p < 0.001. Data represent mean ± SEM.

### Acute stress rapidly and transiently changes the phospho-proteome

Some of the earliest intracellular molecular changes triggered by stress occur at the level of protein phosphorylation (Ai et al., 2017; Caudal et al., 2010; Shen et al., 2004). Phosphorylation changes are highly dynamic and are typically not sustained in the face of protein turnover. However, previous work has shown that after very intense stressors, some phosphorylation changes may persist for one hour (Okamura et al., 2019) or even a day (Musazzi et al., 2017b; Revest et al., 2010). Thus, we first assessed protein phosphorylation in an unbiased way on the level of the proteome. Mice were sacrificed either directly after AS exposure, and at 15, 30 or 45 min after the initiation of stress (Figure 2A). Mice in the control group remained in their home-cage until sacrifice. Because the dorsal and ventral HC (dHC and vHC) are engaged in different brain circuitries and are molecularly very distinct (Fanselow and Dong, 2010; Floriou-Servou et al., 2018), we dissected them separately and performed label-free quantitative phosphorylation analysis by liquid chromatography-tandem mass spectrometry (LC-MS/MS). We used 5-8 mice per group (see Figure 2A), and we treated each mouse as an independent biological sample. Overall, we were able to quantify 16302 distinct modified peptide sequences in our phospho-enriched samples, of which 10132 (62%) were phosphopeptides. For all analyses we used FDR adjusted p-values < 0.05 to determine significance. Immediately after stress, we detected 847 significantly altered modified peptides (794 phosphopeptides; ~93%) in the dHC, and 269 in the vHC (253 phosphopeptides; 94%) (Figure 2B,C). 15 min after stress, this number dropped to 206 modified peptides in the dHC (188 phosphopeptides; 91%), and 94 modified peptides in the vHC (86 phosphopeptides; 91%). At the 30 and 45 min timepoints, we could not detect significant changes anymore, indicating that protein phosphorylation changes after AS occur rapidly and seem to be tightly regulated and short-lived in the hippocampus. Significantly altered peptides were strongly enriched for phosphopeptides (fisher exact test; 6min dHC: p = 1.17 e-15, odds ratio = 1.5, 6min vHC: p = 3.4 e-6, odds ratio = 1.51), and phosphopeptides demonstrated stronger fold-changes compared to non-phosphorylated peptides (Figure S2C). To ensure that the observed phosphorylation changes are not due to overall changes in the proteome (protein abundance), we performed a quantitative proteomic measurement of reference samples (not enriched for phosphopeptides), which revealed no significant changes 6 min after stress (Figure S2B). Importantly, the protein phosphorylation changes were correlated between the 6 and 15 min timepoints (dHC: R^2^ = 0.35, vHC R^2^ = 0.39), as well as between the 15 and 30 minute timepoints (dHC: R^2^= 0.35, vHC R^2^ = 0.55), suggesting that most of the phosphorylation events belong to one big phosphorylation wave that unfolds over time (Figure 2E). In line with known molecular differences between dHC and vHC (Cembrowski et al., 2016a, 2016b; Dong et al., 2009; Floriou-Servou et al., 2018), we observed baseline differences in protein phosphorylation between dHC and vHC (Figure S2D). Despite these baseline differences, there was a strong correlation between the stress-induced protein phosphorylation changes in the dHC and vHC, at both the 6 min (R^2^= 0.42) and 15 min (R^2^= 0.48) timepoints (Figure 2D). This suggests that even though more significantly altered phosphorylations can be observed in the dHC, similar and robust phosphorylation cascades are triggered throughout the longitudinal axis of the HC. Using *K*-means clustering, we grouped the significantly modified peptides in 5 distinct temporal profiles (Figure 2F), and assigned their corresponding proteins to gene ontology (GO) terms. Most modified proteins were related to dendritic morphology and development, calcium signaling and synaptic function, AMPA-receptor regulation and neurotransmitter release. This all points to stress-induced activation of neuronal activity in the HC. Across all swim responsive phosphoproteins we found enrichment for a number of pathways, including calcium signaling (Figure S2A), a pathway well-known to react to neuronal activation (Wiegert and Bading, 2011). A domain enrichment analysis revealed that significantly altered phosphopeptides are overrepresented in two InterPro (Blum et al., 2021) protein domains, the Protein Kinase C and SAPAP (also known as Dlgap) family of proteins (Figure S2E), the latter is associated with synaptic function and is implicated in multiple neuropsychiatric diseases (Rasmussen et al., 2017; Welch et al., 2007).

**Figure 2:**
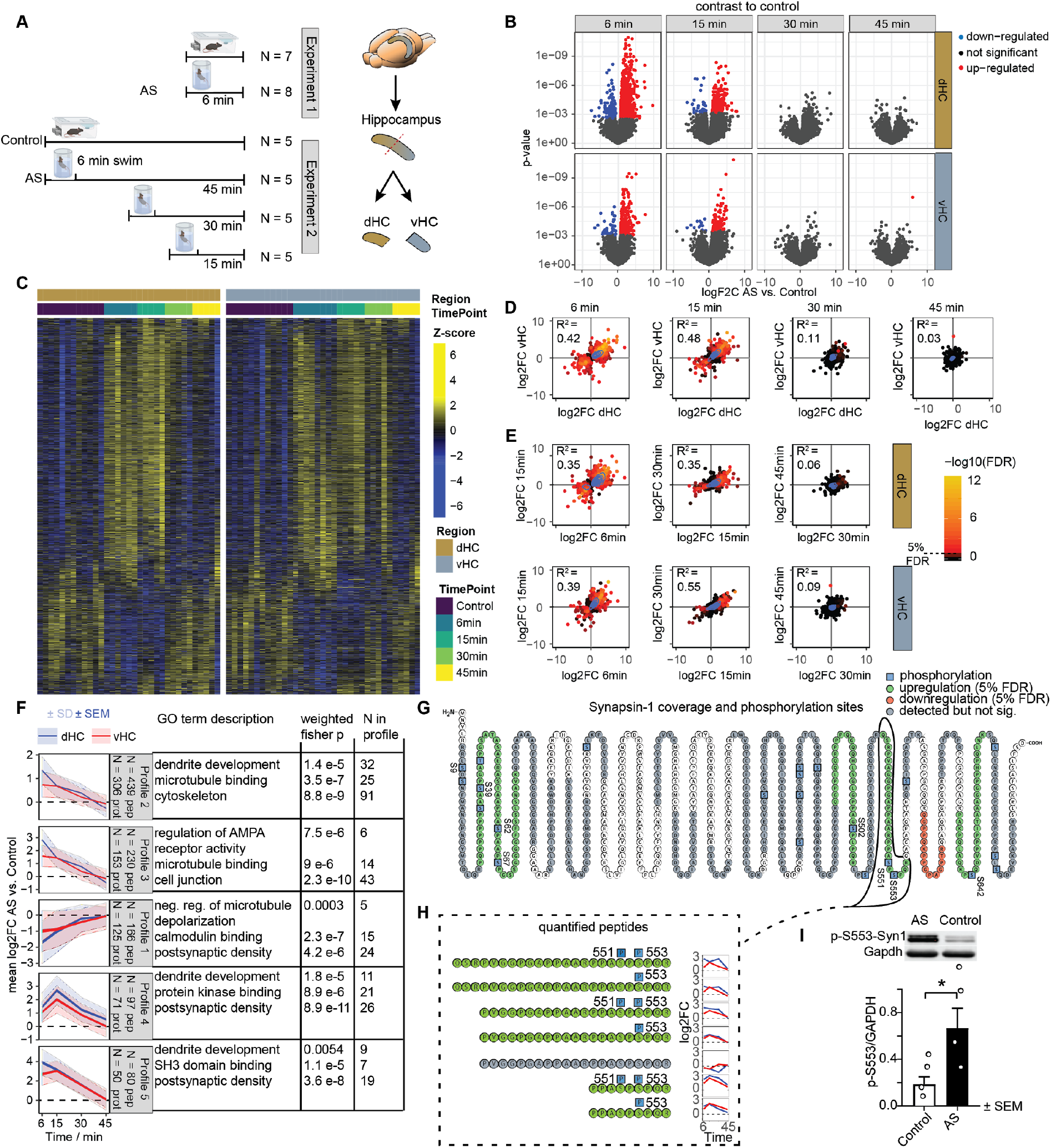
Rapid, short-lived effects of acute stress (AS) on the phosphoproteome of the dHC and vHC. **A)** Experimental design and tissue collection approach. **B)** Volcano plots showing the estimated log2 fold changes and statistical test results for modified peptides immediately (Exp. 1) or shortly (Exp. 2) after exposure to AS. Red and blue dots represent significant changes within 5% FDR relative to non-stressed controls. **C)** Heat maps showing the abundance of significant modified peptides across all samples. **D)** Correlation of modified peptides that are significant at least at one time point, between the dHC and the vHC (all correlations are significant with p < 1.18 e-8) **(E)** Correlation of modified peptides that are significant at least at one time point, between different time points in the dHC and vHC (all correlations are significant with p < 1.11 e-22) **F)** *K*-means clustering of significant modified peptide changes to resolve 5 temporal profiles, and GO term analysis of the corresponding proteins. **G)** Amino acid sequence of Synapsin 1, illustrating overall coverage (grey, green and red), phosphopeptides significantly upregulated (green) or downregulated (red) at any time point after AS and detected phosphosites (blue, PTM probability > 0.75) **H)** Quantification of all detected Synapsin 1 modified peptides spanning sites pS551 and pS553, and their temporal profiles (blue = dHC, red = vHC). **I)** Validation of the upregulation of the phosphorylated site SYN1-pS553 with Western blot in the dHC immediately after stress (Effect size ~300%, t(6) = 2.70, p = 0.0357, unpaired t test, N=4 mice/group). *p < 0.05.

Phospho-proteomics not only provides a uniquely detailed map of the complex and widespread phosphorylation changes detected across proteins, but it also reveals unique details about the phosphorylation changes within individual proteins. To illustrate this, we focused on Synapsin 1 (SYN1), a protein with well-characterized phosphorylation sites. Using a non site-specific antibody, SYN1 has been shown to be phosphorylated at various time points after AS in the HC (Revest et al., 2010), while another study reported increased S9 phosphorylation in the prefrontal cortex up to 24 hours after stress (Musazzi et al., 2017b). By interrogating the whole SYN1 sequence, we detected both upregulated and downregulated phosphopeptides immediately after AS (Figure 2G), with most detected peptides being phosphorylated at serine 553 upon stress (p-S553) (Figure 2H). Indeed, western blot analysis validated a 3-fold increase in the levels of p-S553-SYN1 (Figure 2I) in the dHC immediately after stress. Other significantly upregulated phosphopeptides contained the phosphosites S39, S62, S67, S502, S551, S642, although the phosphorylation at S9 did not pass our FDR cutoff (logFC 6min vHC 1.32, p = 0.005, adj.p = 0.16). Most of these phosphorylation changes were still detectable at 15 min, but diminished at later time points. Together, these data reveal the stunning complexity of the phosphorylation code at the level of a single protein, as well as the breath of the phospho-proteomic changes triggered by AS.

### Stress-induced transcriptomic changes evolve over time

Protein phosphorylation triggers second messenger cascades that lead to changes in gene expression. However, it remains unknown how the stress-induced transcriptomic changes evolve over time, and whether these events unfold similarly in the dHC and vHC. Therefore, we profiled the transcriptome of the dHC and vHC at 45, 90, 120, 180 or 240 min after the initiation of stress using bulk RNA-sequencing (RNA-seq, Figure 3A). We used 7-8 mice per group, and we treated each mouse as an independent biological sample.

**Figure 3:**
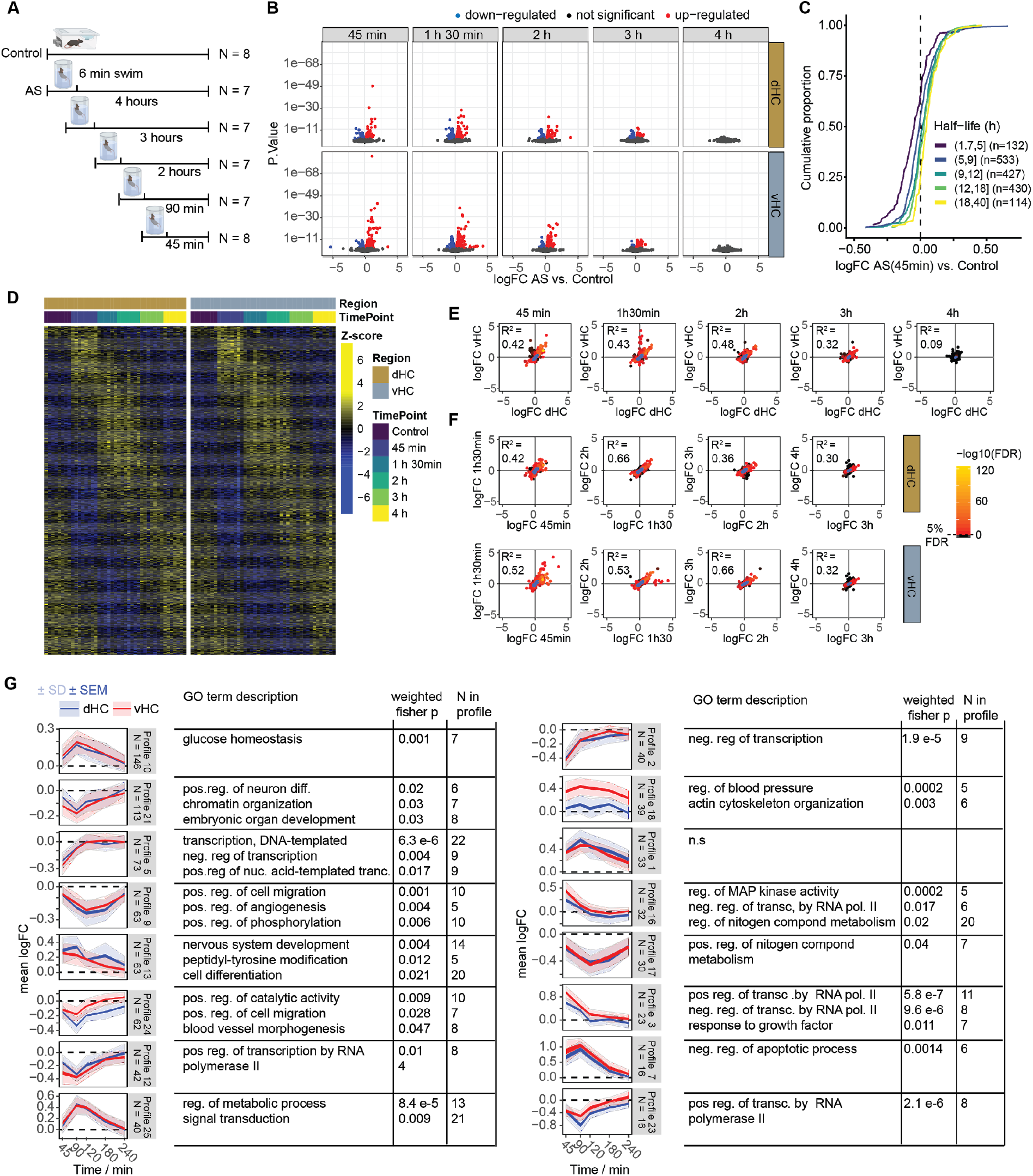
The effects of acute swim (AS) stress on the transcriptome of the dHC and vHC across time. **A)** Experimental design. **B)** Volcano plots showing the log2 fold change and statistical results of RNA transcripts at 5 different timepoints after exposure to AS. Red and blue values represent changes within 5% FDR. (N = 7-8 per group) **C)** Cumulative fold change distributions of genes with different half-lives, consistent with the downregulation being the result of normal mRNA decay. **D)** Heat maps showing expression of all differentially-expressed genes in all samples. **E)** Correlation of transcriptional changes significant in at least one timepoint, between the dHC and the vHC (all correlations are significant with p < 4.8 e-19). **(F)** Correlation of transcriptional changes significant in at least one timepoint, between consecutive timepoints in the dHC and vHC (all correlations are significant with p < 2.5 e-70). **G)** *K*-means clustering of differentially expressed RNA transcripts to resolve temporal profiles, and their GO term description. *K*-means clustering was done with 25 centers, of which the ones with less than 15 transcripts were excluded. Blue indicates expression in dHC, red in vHC. Plots indicate both standard error of the mean (SEM) of logFC in bright color and standard deviation (SD) in faint color.

We observed highly dynamic gene expression changes over time in response to AS. Using 5% FDR-corrected p-values, the highest number of gene expression changes occured at 45 and 90 min after stress, followed by a gradual decline in both dHC and vHC (Figure 3B,D). After 4 hours, no significant changes were detected anymore, suggesting a tight regulation of gene expression over time. We observed a high correlation in the gene expression changes between consecutive timepoints up to 3 hours after stress, indicating that these changes are not random but rather systematic and evolve over time (Figure 3F). Overall, we detected more stress-induced gene expression changes in the vHC than in the dHC, consistent with our previous work (Floriou-Servou et al., 2018). However, we found that significantly changed genes correlated well between the dHC and vHC at all early timepoints (45 min: R^2^= 0.42; 90 min: R^2^= 0.43; 2 hours: R^2^= 0.48) indicating a similar pattern of gene expression changes (Figure 3E) along the longitudinal axis of the HC.

We next clustered the significant genes based on their temporal transcriptional profiles and characterized the clusters in terms of their top enriched GO biological processes (Figure 3G). As expected, genes whose expression peaked at the earliest time point and then rapidly decreased (profiles 3 and 16) were associated with MAPK activity and the regulation of transcription. A second wave of genes peaked at 90 min, and was associated with glucose homeostasis (profile 10) and metabolism (profile 25), while apoptotic programs (profile 7) were negatively regulated. Later timepoints did not appear to show novel distinct transcriptional programs, but rather the gradual return to baseline transcription.

In parallel to these waves of transcription, an equally large number of genes rapidly decreased in expression. This could be the consequence of active RNA degradation, for instance mediated by micro-RNAs, or due to rerouting of the transcriptional machinery and normal RNA decay. To investigate the former, we performed small RNA sequencing of both hippocampal regions 45 min after stress. While quality control was successful (Figures S3A,B), we found no significant change in miRNA expression, nor any significant miRNA target enrichment in the transcriptional signature (Figure S3C,D). To test the hypothesis that the rapid down-regulation was instead the result of normal mRNA decay, we plotted the fold changes for genes of different half-lives, as estimated in unperturbed mouse fibroblasts (Schwanhäusser et al., 2011). This revealed that genes with shorter half lives were more strongly downregulated (Figure 3C). In addition, downregulated genes have a median half-life of 4.5 hours in the dish, with an expected passive downregulation of 11% after 45 min, which is consistent with the observed median downregulation of 12% across the same genes 45 min after AS. Together, these results suggest that stress strongly drives a first, regulatory wave of transcription, followed by a second wave that reestablishes metabolic homeostasis. We speculate that this major transcriptional activity occurs at the expense of other genes which decrease largely due to normal RNA decay.

As the experiments reported thus far were performed in male mice, and there have been reports of sex-differences in the stress response (Bale and Epperson, 2015; Bangasser and Wicks, 2017), we compared the transcriptional response to stress in males and females (see Figure S4A for experimental design). We found that both sexes reacted remarkably similar to AS. We faithfully reproduced the strong transcriptional response observed in males at the 45 minute timepoint with no significant interaction between sex and treatment, and we again confirmed that all gene expression changes disappeared within four hours (Figure S4C,D). We did detect sex differences in gene expression at baseline (Figure S4D-F), however noted that these are mainly x- and y-chromosomal genes. We also addressed whether the transcriptional response to AS is similar in the left and right hemispheres, as lateralization effects had previously been reported (Igloi et al., 2010; Kohl et al., 2011; Miller et al., 2018). A direct comparison of RNAseq data from left vs. right hemisphere in the same cohort of mice 45 min after AS (Figure S4B) shows that the molecular response in both hemispheres is extremely similar, with no significant interaction between hemisphere and treatment (Figure S4G,H).

### Active nuclear transcription and transcription factor activity after stress

In the process of mRNA maturation, RNA is first transcribed in the nucleus as unprocessed pre-RNA, before being spliced relatively quickly and transported out of the nucleus for translation (Alpert et al., 2017). Therefore, we used unprocessed transcripts as a proxy for actively ongoing nuclear transcription. By independently analyzing processed and unprocessed transcripts, we found that altered nuclear transcription mainly occurs early, at 45 min after stress, and to a far lesser extent at the 90 min and 2 hour timepoints (Figure 4A). In contrast, processed mRNAs persist at later timepoints and resolve more slowly (Figure 4A). A PCA analysis demonstrates that the unprocessed response is shorter in duration and relaxes much faster along the first principal component in contrast to the processed response (Figure 4B). Also within the profile clusters described above, unprocessed expression changes precede and predict expression of processed transcripts (Figure 4C). The independent analysis of unprocessed and processed transcripts further enables more accurate estimates of transcription factor (TF) activity. Many TFs have complex modes of action (e.g. where they are activated through PTMs, altered shuttling to the nucleus and/or binding to other cofactors), which cannot be inferred from their RNA levels. In addition, our phosphoproteomic data did not detect many TFs, preventing us from directly linking phosphorylated peptides to TF activity. We therefore opted to infer TF activity from the relative expression of their targets’ unprocessed transcripts based on curated regulons (Garcia-Alonso et al., 2019). We identified TFs for which our data suggest increased activity mainly at the early 45 minute time point (Figure 4D,E and Figure S5A). One example is *Creb1*, which is activated through calcium signaling (Sakamoto et al., 2011), a pathway for which we saw many of its components (i.e. signaling proteins such as CAMKs and PKC) altered in our phospho-proteomics data (Figure S2A,E). To identify distinct transcriptional programs underlying the observed expression changes, we clustered differentially-expressed genes based on their temporal expression profile, and their inferred regulation by the TFs. This revealed transcriptional programs with significant enrichment for very distinct biological processes, which again include glutamatergic and synaptic networks (Figure 4E). Overall, AS triggers active transcription very rapidly, and during the subsequent 4 hour period, waves of gene expression unfold and fully return back to baseline.

**Figure 4:**
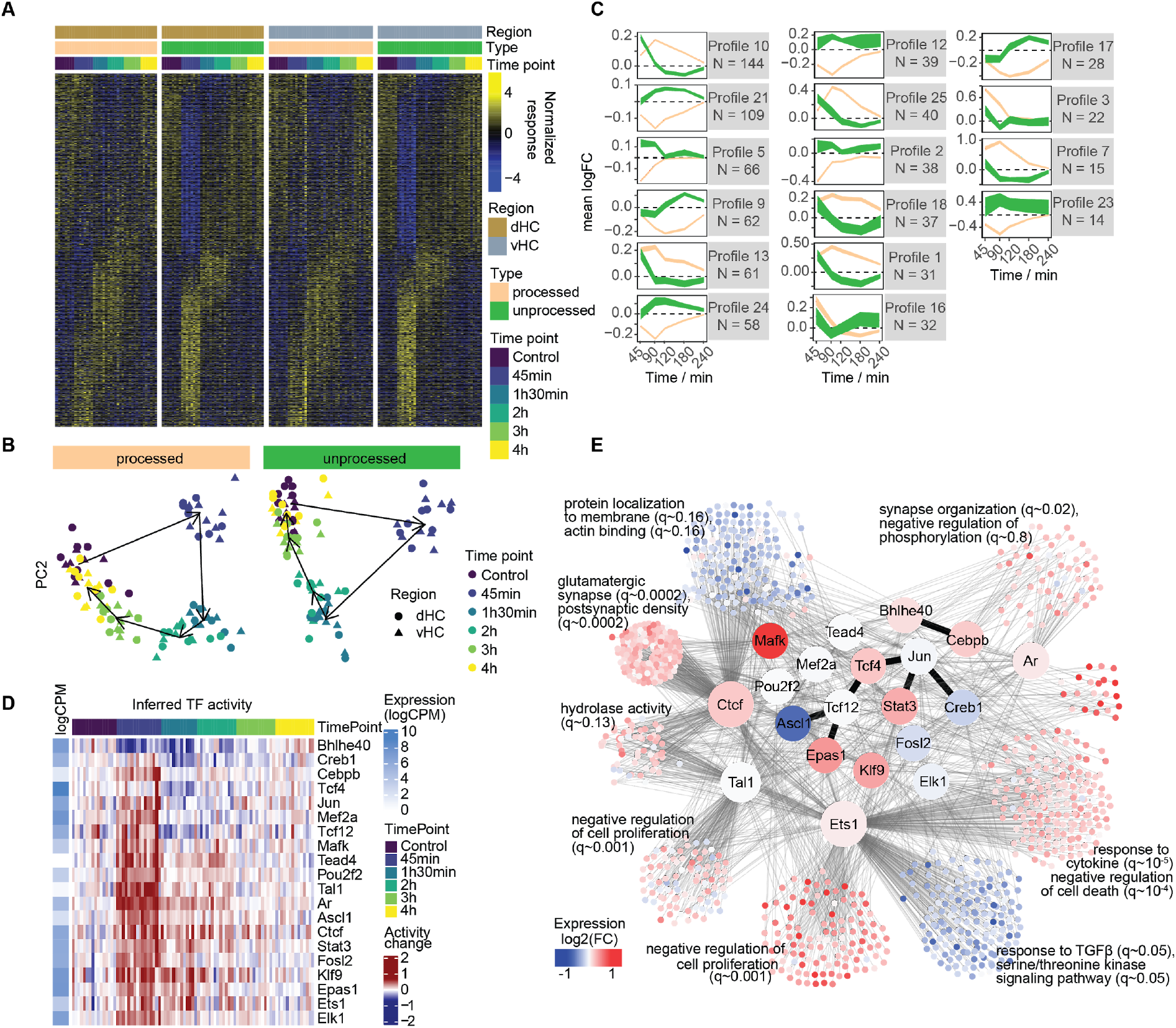
Active nuclear transcription after acute stress. **A)** heatmap showing unprocessed (actively transcribed) vs processed transcripts of differentially expressed genes. **B)** PCA plot of unprocessed and processed transcripts across differentially-expressed genes. **C)** Profiles from Figure 3 resolved by unprocessed (green) and processed (brown) transcripts. **D)** Relative activity of the most significant transcription factors (TF), inferred from the expression of their targets. **E)** Network representation of the curated interactions between differentially-expressed genes and the TFs found to be differentially active. Colours indicate the direction of differential expression at 45 min. Thin lines represent transcriptional interactions, while dark, thick lines represent protein-protein interactions between the factors. For each cluster, the top GO enrichments are shown.

### Specific stress-induced transcriptomic changes in different cell-types

To test whether stress-induced transcriptional changes occur in specific cell types, we performed single nucleus RNA sequencing (snRNA-seq) after AS. Because nuclear RNA was sequenced, we focused on the 45 minute timepoint, when nuclear transcription is most active. We observed a total of 5292 QC-filtered nuclei in four independent hippocampal samples from 2 control mice and 2 stress-exposed mice. A combination of clustering and reference-based assignment produced distinct cell clusters linked to specific cell-types or families (Figure 5A). We first inquired in which cell types differentially expressed transcripts from the bulk sequencing are most abundantly expressed. We found that overall these genes were mainly expressed in neurons, astrocytes, oligodendrocytes and vascular cells, but less in microglia and oligodendrocyte precursor cells (OPC) (Figure 5B). Most upregulated genes from bulk sequencing show the highest expression in astrocytes. To assess the extent to which snRNA-seq results mirror bulk sequencing results in the response to AS, we analyzed how well their log-fold-changes correlated. Neurons and astrocytes showed the best correlation with unprocessed bulk mRNA (Figure 5C). However, since these correlations could be confounded by the relative abundance of the different cell types, we confirmed these results with an alternative assignment-based approach (see methods). In line with the correlations, we found that the majority of expression changes (49%) were assigned to glial cells, whereas a smaller number of gene expression changes were assigned to neurons (24%) and vascular cells (18%) (Figure 5D). Many genes showed strong, cell-type specific regulation (Figure 5E). *Apold1*, a known endothelial gene (Regard, 2004) that is strongly upregulated by AS (Floriou-Servou et al., 2018; Roszkowski et al., 2016), indeed showed a strong stress-induced increase only in vascular cells. In contrast, we found genes such as *Dio2* and *Map3k19*, which seem to be very selective for astrocytes. Overall, these results highlight that - beyond neurons - glial cells and vascular cells contribute strongly to the overall transcriptional response to stress, and that this response is also highly cell-type specific.

**Figure 5:**
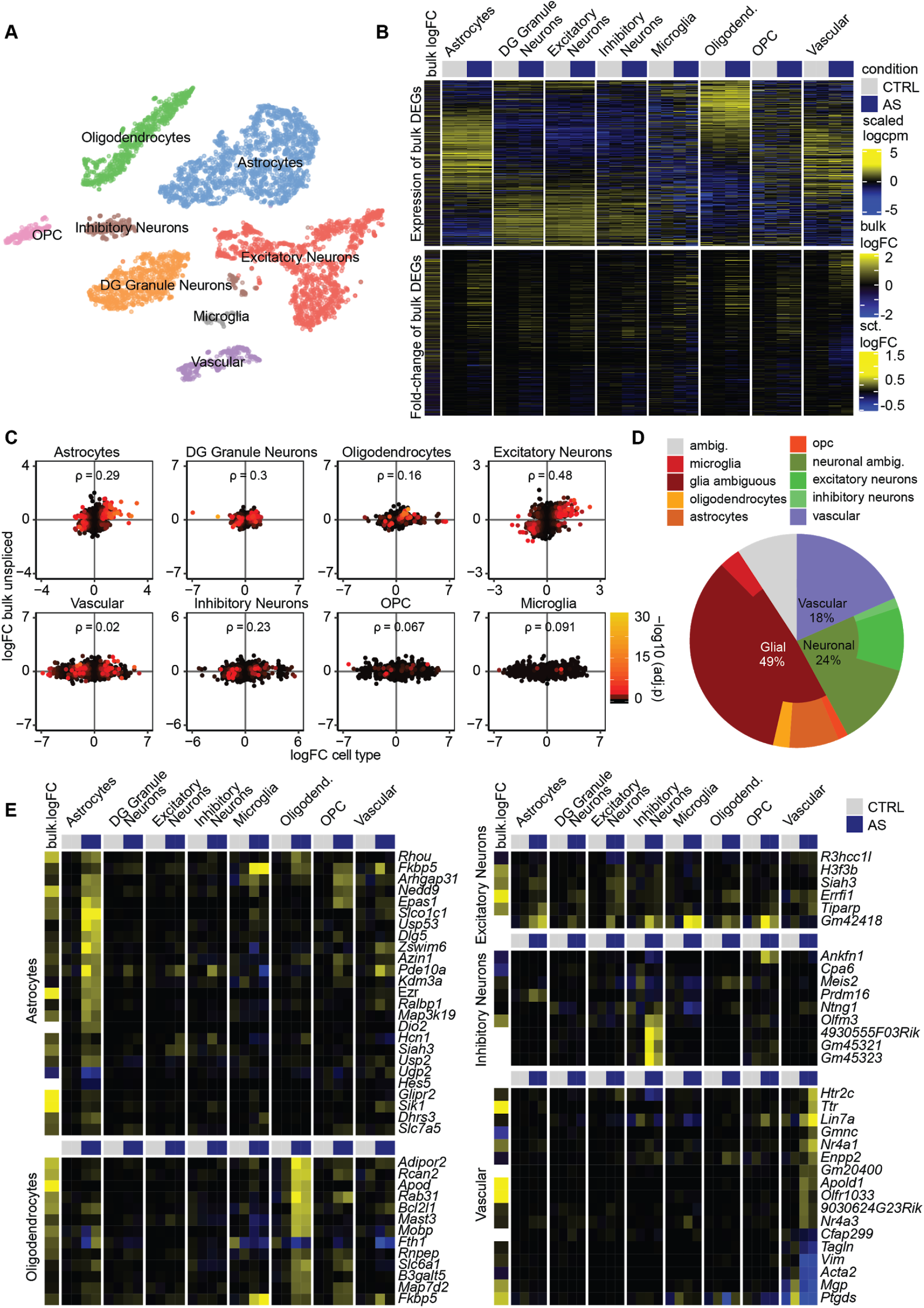
Single nucleus RNA sequencing 45 minute after acute stress (AS). **A)** t-SNE plot of single-nuclei transcriptomes labeled by cell-type. **B)** *Top:* Heatmap indicating overall expression of stress responsive genes in logcpm (from bulk sequencing) in snRNA cell-types. *Bottom:* Heatmap indicating logFC after AS of these genes in the snRNA data. **C)** Correlations between logFCs of DEGs in bulk sequencing (45 minutes unprocessed) and logFC of snRNA data within cell-types (ρ = Spearman’s rank correlation coefficient). Color indicates adjusted p-value in snRNA data. **D)** Decomposition analysis highlights which cell-types best explain DEGs from the bulk sequencing in the snRNA data. **E)** Top gene candidates from the differential expression analysis in snRNA. Each heatmap shows top candidates of a selected cell-type and their expression in single-nuclei aggregated by cell type.

### Stress-induced effects on the translatome

Our transcriptomic analyses have revealed widespread and profound changes in gene expression across time and in different cell types. To determine how many of these transcripts will then become actively translated, we measured actively translated RNA using translating ribosome affinity purification (TRAP) followed by RNA sequencing (TRAP-seq) (Heiman et al., 2014; Picelli et al., 2014) (Figure 6A). To target actively translated RNA in all brain cells, we crossed CMV-Cre mice (Schwenk et al., 1995) with a floxed-nuTRAP mouse line (Roh et al., 2017). CMV-nuTRAP mice ubiquitously express a ribosomal green fluorescent protein (eGFP) tag fused with the ribosomal subunit L10a (Figure 6B) and an mCherry tag fused with the nuclear pore protein RanGAP1. This experiment was performed only in females, because the breeding generated few double-transgenic males. To target actively translated RNA specifically in excitatory neurons, we crossed CaMKIIa-Cre mice with floxed-nuTRAP mice (Figure 6C). To target inhibitory neurons, we used the vGAT::bacTRAP mouse line (Das Gupta et al., 2021), expressing the same eGFP ribosomal tag under the vesicular GABA transporter (vGAT, Slc32a1) promoter (Figure 6D). CaMKIIa-NuTRAP and vGAT::bacTRAP experiments were conducted in males.

**Figure 6:**
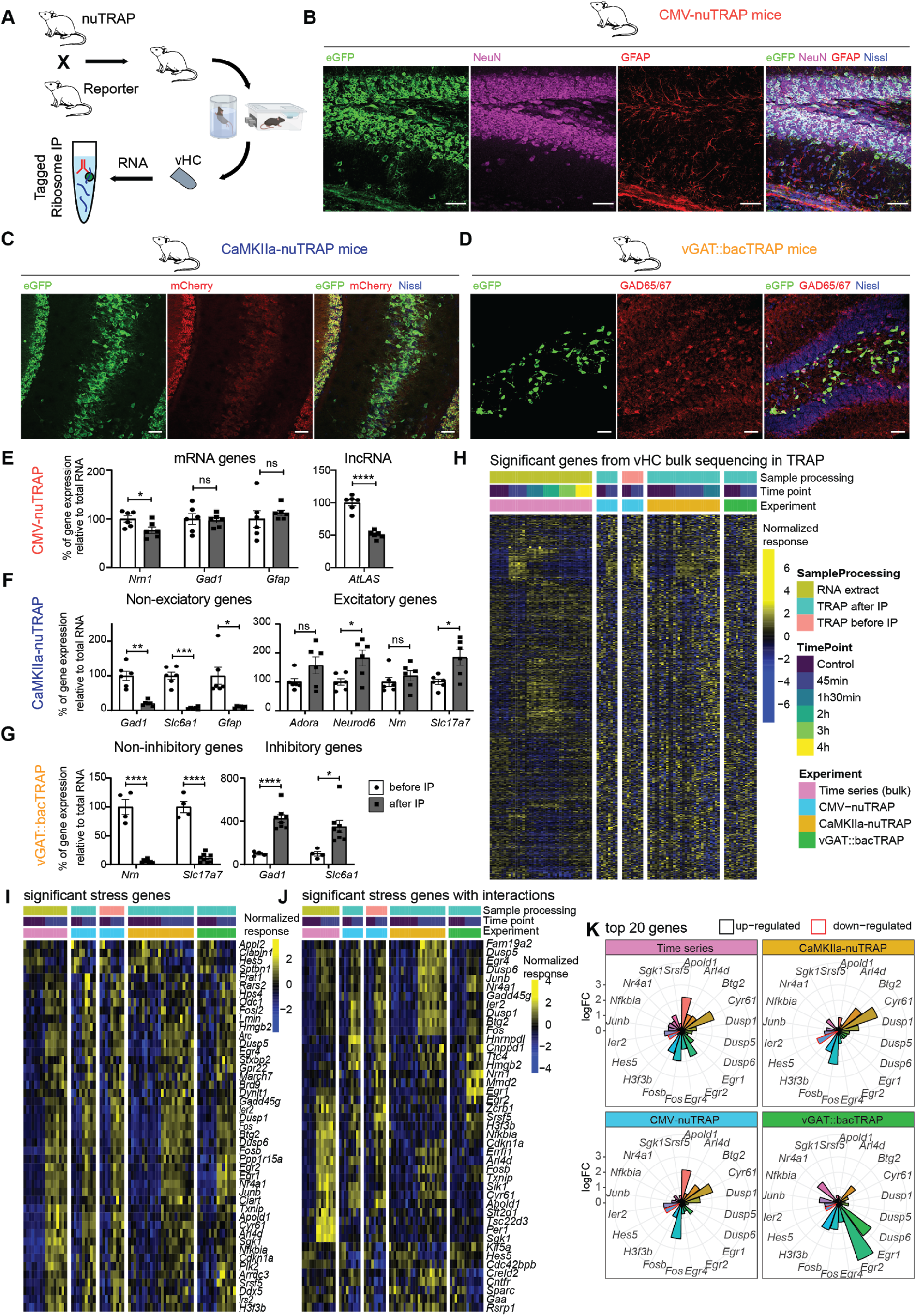
Acute stress (AS) effects on the whole translatome, and the translatome of excitatory and inhibitory neurons of the vHC. **A)** Schematic of the TRAP approach, showing that nuTRAP mice are crossed with cell-type specific reporter lines. Then, animals are subjected to AS, subsequently dHC and vHC are dissected and samples are enriched for tagged ribosomes using the TRAP protocol. **B)** Microscopy images showing co-localization of eGFP with NeuN and GFAP in the hilus and DG of a CMV-nuTRAP mouse. **C)** Microscopy images showing co-localization of eGFP fused with L10a, and mCherry fused with RanGAP1 in the ventral hilus, CA3 and DG of a CaMKIIa-nuTRAP mouse. **D)** Microscopy images showing co-localization of eGFP fused with L10a, and GAD65/67 in the dorsal hilus and DG of a vGAT::bacTRAP mouse. **E-G)** RT-qPCR validation of TRAP enrichment. Individual values represent dHC and vHC samples from 3 biological replicates. **E)** Genes primarily expressed in excitatory neurons, inhibitory neurons and glial cells are present both before (total RNA) and after (bound RNA) the IP, while the long non-coding RNA (lncRNA) *AtLAS* is reduced in the bound RNA of hippocampal samples from CMV-nuTRAP mice (N = 3). RNA levels were normalized to *Hprt*. Asterisks indicate p-values generated by independent unpaired t-tests (*Nrn1* t(10) = 2.486, p = 0.0322; *Gad1* t(10) = 0.2409, p = 0.8145; *Gfap* t(10) = 0.683, p = 0.5101); *AtLAS* t(10) = 9.134, p < 0.0001. Data represent mean ± SEM. Scale bars 50 μm. **F)** Depletion of genes not expressed in excitatory neurons, and enrichment of genes primarily expressed in excitatory neurons after IP (bound RNA) compared to before (total RNA), in hippocampal samples from CamKIIa-nuTRAP mice (N = 3). RNA levels were normalized to *Tubd1*. Asterisks indicate p-values generated by independent paired t-tests (*Gad1* t(5) = 6.65, p = 0.0012; *Slc6a1* t(5) = 9.00, p = 0.0003; *Gfap* t(5) = 3.69, p = 0.0141; *Adora1* t(5) = 2.20, p = 0.0794; *Neurod6* t(5) = 3.32, p = 0.0210; *Nrn1* t(5) = 1.15, p = 0.3012; *Slc17a7* t(5) = 2.87, p = 0.0350). **G)** Depletion of genes not expressed in inhibitory neurons and enrichment of genes primarily expressed in inhibitory neurons after IP (bound RNA) compared to before (total RNA), in hippocampal samples from vGAT::bacTRAP mice (N = 4). RNA levels were normalized to *Gapdh*. Asterisks indicate p-values generated by independent unpaired t-tests (*Nrn1* t(10) = 10.49, p< 0.0001; *Slc17a7* t(10) = 11.67, p < 0.0001; *Gad1* t(10) = 6.23, p < 0.0001; *Slc6a1* t(10) = 3.01, p = 0.0131). **H)** Heatmap illustrating the normalized response of significant stress genes (from the bulk sequencing) in the time-series data vs. the TRAP samples in the vHC. (CMV-nuTRAP N = 4 x 5 (pre-IP), 4 x 5 (post-IP); CaMKIIa-nuTRAP N = 12(ctrl) x 12(45min) x 7(90min); vGAT::bacTRAP N = 7 x 7) **I)** Significant genes that are AS responsive in all experiments (time series, CMV-nuTRAP, vGAT::bacTRAP and CaMKIIa-nuTRAP) in the vHC. **J)** Significant genes when employing a model that includes interaction terms in the vHC. **K)** logFC 45 min after AS across datasets of the top 20 genes in the vHC.*p < 0.05, **p < 0.01, ***p < 0.001, ****p < 0.0001. Data represent mean ± SEM. Scale bars 50 μm.

To validate the TRAP method, we performed RT-PCR to compare the expression of mRNAs and long non-coding RNAs in CMV-nuTRAP pre and post immunopurification (IP) fractions. We found a significant depletion of lncRNAs, whereas mRNAs were not changed (Figure 6E). Further, we confirmed expression of inhibitory and excitatory genes in the CaMKIIa-nuTRAP and vGAT::bacTRAP samples before IP (total RNA) and afterwards (bound RNA). As expected, inhibitory and glial genes were depleted, and excitatory genes were largely enriched in the bound RNA fraction of CaMKIIa-nuTRAP HC samples (Figure 6F). Conversely, excitatory genes were depleted and inhibitory genes were enriched in the bound RNA fraction of vGAT::bacTRAP tissue (Figure 6G). However, the enrichment of excitatory genes seems to be less efficient in CaMKIIa-nuTRAP samples. This is likely due to several reasons: (1) excitatory neurons are the majority of neurons in HC, and hence it is harder to show enrichment of this abundant cell type. (2) These genes might not be exclusively expressed in excitatory neurons. (3) Expression of these genes is highly variable in both pyramidal and granular cells along the dorsoventral axis (Cembrowski and Spruston, 2019; Cembrowski et al., 2016a, 2016b), hence the higher variability lowers the statistical power to show significant enrichment. To confirm the specificity of our TRAP lines we used immunohistochemistry. eGFP was expressed in both neurons and glial cells in CMV-nuTRAP mice (Figure 6B). Further, we confirmed eGFP and mCherry expression in pyramidal cells in the HC of CaMKIIa-nuTRAP mice (Figure 6C), and eGFP expression and co-localization with glutamate decarboxylase 65/67 (GAD65/67) in the HC of vGAT::bacTRAP mice (Figure 6D).

Next, we subjected CMV-nuTRAP, CaMKIIa-nuTRAP and vGAT::bacTRAP mice to AS, and collected the vHC 45 min after onset of stress for TRAP-seq. We used 4-7 mice per group, and we treated each mouse as an independent biological sample (see Figure S6A for full overview of all experimental designs). TRAP-seq analysis revealed that only a small number of genes were significantly changed in either the whole translatome (in vHC), or in the translatome of excitatory or inhibitory neurons in the vHC (Figure S6C) and dHC (Figure S6B). This suggests that only few of the genes that get transcribed also become actively translated, reminiscent of translation inhibition during the integrated stress response (Advani and Ivanov, 2019). To ensure that we did not miss translational changes because transcription might not yet occur 45 min after stress, we repeated the Camk2a-nuTRAP experiment 90 minutes after stress exposure. Again, the translatomic effects closely mimicked the results obtained at the 45 minute time point (Figure S6C), demonstrating the reproducibility of our approach and ruling out that the low number of translated mRNAs is associated with the timing of our analysis. Despite the high number of biological replicates used for our analyses, we noticed that variability in TRAP samples was generally higher than in transcriptomic analyses (figure S6D). This is in line with a recent reanalysis of TRAP-seq data (von Ziegler et al., 2020) and suggests that statistical power of TRAP-seq analysis is lower than that of bulk RNAseq. To increase statistical power, we performed a meta analysis which included data from both the transcriptome time series assay, as well as all TRAP-seq samples. We used surrogate variable analysis (SVA) (Leek et al., 2012) to remove technical variability across batches of samples. We visualized the TRAP-seq data for those genes that were differentially regulated by AS in the bulk sequencing experiment (Figure 6H). We found that in the vHC a subset of the genes altered at 45 min in the transcriptome are also altered in the translatome. Notably, this was mainly restricted to up-regulated genes. Adjusting for baseline differences across datasets and differences in response strength, we identified genes that demonstrated consistent and significant changes across datasets 45 min after stress (Figure 6I). These bona-fide stress-induced genes include many well-known immediate early genes such as *Fos, Arc, Sgk1, Egr1, Dusp1* and *Apold1*, indicating that active translation of these genes coincides with increased transcript availability. To detect different response strengths or response directions between datasets, we further employed a more complex statistical model with interaction terms (see methods) accounting for these differences. We found numerous genes that were significantly altered (Figure 6J), most of which also had significant interactions (Figure S6E). One thing this analysis highlights is the remarkable cell-type specificity in the translational response to AS. Translation for genes such as *Egr1, Egr2* and *Nr4a1 was* increased in inhibitory neurons (Figure 6K), while genes such as *Cyr61* and *Btg2* are translated more intensively in excitatory neurons. *Apold1*, an endothelial immediate early gene, is up-regulated much less in inhibitory and excitatory neurons compared to the CMV-nuTRAP and bulk sequencing. Overall, these results demonstrate that different transcriptional and translational programs play out across cell types, and that the initially strong and widespread molecular response is suppressed towards protein translation.

### The proteome in different hippocampal subregions after acute stress

The finding that stress-induced translational changes are much more modest than the phospho-proteomic and transcriptional changes, supports previous findings that hardly any stress-induced proteomic changes can be detected in the dHC and vHC 24 hours after AS (Floriou-Servou et al., 2018). We reasoned that stronger proteomic changes might occur right after the transcriptional response has played out 4-hours after stress. While single cell proteomic analyses are not technically possible, proteomic profiles differ dramatically between dHC and vHC (Floriou-Servou et al., 2018), and also between CA1 and CA3 subregions (von Ziegler et al., 2018). To control for this variability and increase analytical power, we decided to assess proteomic changes 4 hours after the initiation of stress separately in the CA1, CA3 and DG areas of both the dHC and vHC (i.e. 6 regions per mouse), using data-independent label-free proteomics (Figure 7A).

**Figure 7:**
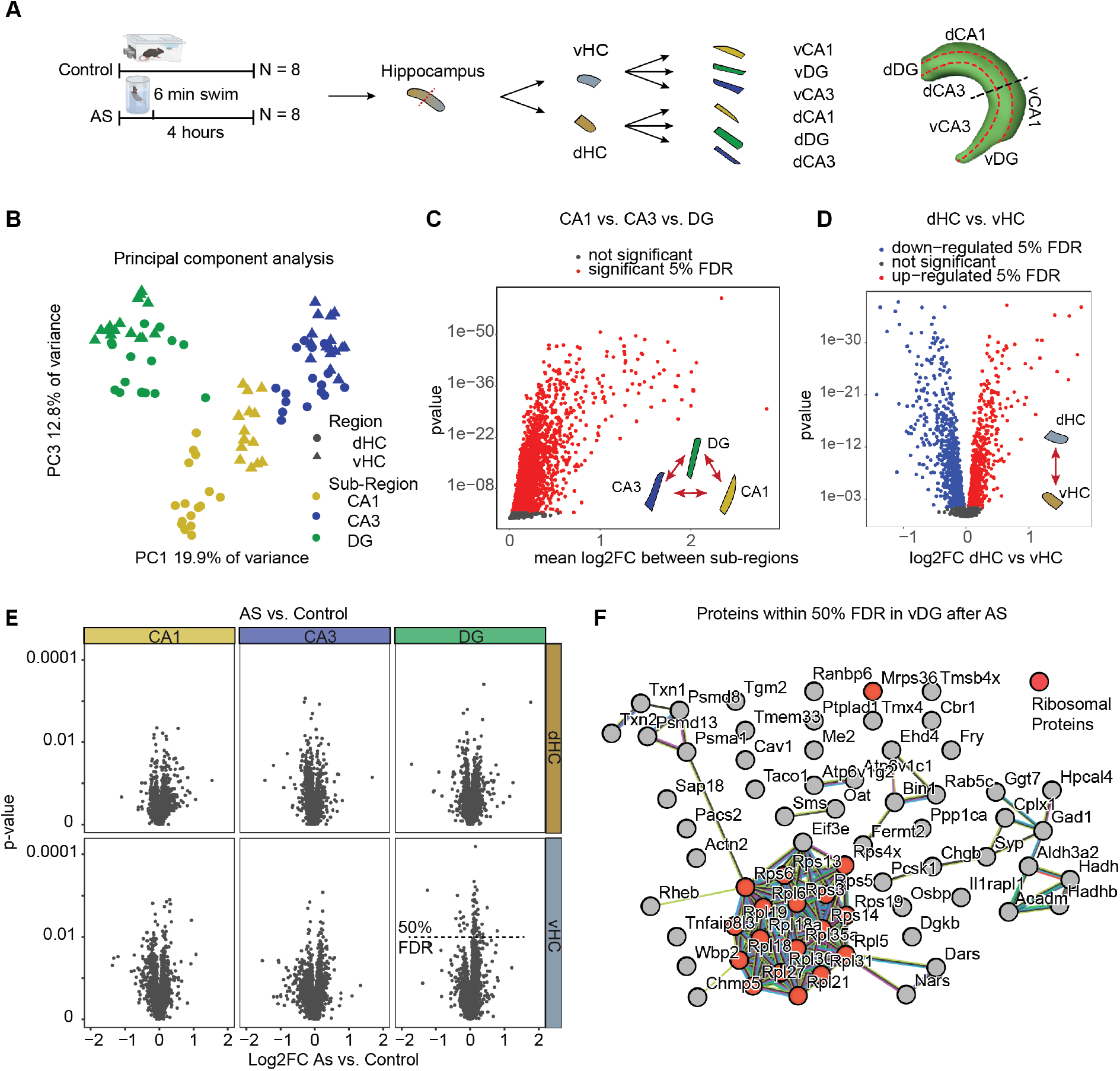
Acute stress (AS) effects on the proteome of CA1, CA3 and DG in the dHC and vHC. **A)** Experimental design and tissue collection approach. **B)** Principal component analysis of all samples. The first component resolves sub-regional differences, whereas the 3rd further resolves dorsal-ventral differences. We did not find any component that resolves stress. **C)** Statistical results for ANOVA subregion * region within animals (omitting stress vs homecage as variable). Many proteins have significant (red, within 5% FDR) sub-regional expression differences at base-line. **D)** Same analysis as in C. Many proteins show significant dorso-ventral expression differences at base-line (red, up-regulated in dHC 5% FDR, blue down-regulated in dHC 5% FDR). **E)** Volcano plots showing the log2 fold change and statistical results of the 2 group comparisons homecage vs swim of proteins 4 hours after a 6 minute exposure to AS in all regions and sub-regions. No proteins are within the 5% FDR, some proteins reach a 50% FDR only in vDG (indicated by dashed line). **F)** protein-protein interaction network of stress sensitive proteins in the vDG (50% FDR).

A principal component analysis showed that both sub-regional differences as well as dorso-ventral differences could be resolved on the proteomic level, confirming strong baseline differences between proteomic profiles of all regions (Figure 7B). Statistical analyses revealed that most proteins (N = 2536 of 3039) had profound sub-regional expression differences between CA1, CA3 and DG (Figure 7C). Further, we found that roughly half of all proteins (N = 1762) had different baseline expression levels between dHC and vHC (Figure 7D). Overall, this demonstrates that strong sub-regional and dorsal-ventral baseline differences in protein expression are present, which confirms and extends our previous proteomic work (Floriou-Servou et al., 2018; von Ziegler et al., 2018), and is in line with the well-characterized transcriptional heterogeneity across hippocampal subregions (Cembrowski and Spruston, 2019; Cembrowski et al., 2016a, 2016b, 2018; Erwin et al., 2020).

We did not detect any significant stress-induced changes in protein expression in any of the subregions (Figure 7E, FDR < 0.05). Only by increasing the significance cutoff to a FDR of 0.5 could we detect some significant changes, although exclusively in the vDG. Further, these proteins were strongly enriched for ribosomal proteins (Figure 7F). To leverage against a slightly higher biological coefficient of variation observed with proteomics compared to transcriptomics (Figure S7A-C), we used larger group sizes for the proteomic analyses. This resulted in similar power for both methods (Figure S7D), suggesting that the absence of stress-induced proteomic changes is not a result of low statistical power.

## Discussion

Here we present the first in-depth, multiomic characterization of the AS response in the mouse HC. Our results reveal a pronounced molecular stress response, which starts with an immediate, broad wave of protein phosphorylation that resolves rapidly between 15 to 30 min after the onset of stress. This is followed by strong active nuclear transcription that quickly resolves between 45 to 90 min after stress. Waves of gene expression peak between 45 to 90 min and completely resolve by 3 to 4 hours. A substantial reduction of mRNAs also occurs during this period, which appears to be largely due to normal mRNA decay. Using snRNA-seq we show that many genes have remarkable cell-type specificity, and glial cells contribute most to the transcriptional bulk sequencing signal. Surprisingly, only a subset of differentially expressed genes become actively translated, and again the translational response varies markedly between different cell-types. At the level of the proteome, we detect profound baseline differences between subregions of the HC, but hardly any stress-induced changes. Since stress can differentially impact dHC and vHC (Floriou-Servou et al., 2018; Maggio and Segal, 2007; Segal et al., 2010), we conducted all our analyses in both fractions of the HC. Although we find extensive baseline differences between vHC and dHC on all molecular levels, we observe a rather homogeneous stress-response in both regions. Overall, molecular changes can differ in magnitude between these regions, however changes are usually strongly correlated, suggesting coordinated molecular responses that unfold throughout the HC. This is in line with a recent meta analysis, which revealed that the molecular response to AS is rather uniform across the dorsal-ventral axis (Floriou-Servou et al., 2021). Importantly, the transcriptional response we observe after AS is remarkably similar to the changes described after novelty stress or restraint stress (Floriou-Servou et al., 2021), suggesting our results are also relevant for different stress modalities.

### Stress effects on the phosphoproteome

To our knowledge, this is the first description of phospho-proteomic changes after AS. Our employed methods do not allow separate analysis of different cell types, which remains a major challenge in neuroproteomics (Wilson and Nairn, 2018). However, overrepresentation analysis suggests that a large part of the response occurs in neurons. In agreement with previous literature, we find widespread stress-induced changes in pathways related to dendritic morphology, synaptic function and plasticity, like the calcium-dependent neuronal MAPK/ERK pathway (Blume et al., 2008; Inda et al., 2017; Refojo et al., 2005; Shinohara et al., 2018). We observed increased phosphorylation of MAPK2, P38 MAPK, CAMKII, and SYN1, which are linked to synaptic plasticity (Ahmed, 2006; Cesca et al., 2010; Shen et al., 2004). Notably, SYN1 can stay phosphorylated up to 24 hours after an intense, inescapable 40 minute footshock stressor (Musazzi et al., 2017b). The fact that we did not detect any lasting effects beyond 30 min could indicate that the response to brief AS can be effectively terminated, but that more intense stressors may dysregulate the system, a hypothesis that needs to be investigated. Beyond known pathways, we identify large numbers of phosphorylated proteins that - to our knowledge - have never been reported to change after an AS exposure. Notable candidates include the proteins SHISHA6, SHISHA7, SHANK1, SHANK2 and SHANK3, which are related to postsynaptic density and plasticity.

### Stress effects on gene transcription

Although previous work has assessed the transcriptomic changes after AS (for review see (Floriou-Servou et al., 2021)), our results describe the temporal dynamics of the transcriptional response from start to end. We find groups of genes with specific temporal response profiles. Most gene clusters that peak early (45 min) after stress are involved in the regulation of transcription and intracellular signaling cascades (e.g. regulation of MAPK activity). This is in line with earlier reports that the MAPK/ERK pathway regulates transcription by inducing *Fos* and *Egr1* in the HC (Chandramohan et al., 2008; Gutièrrez-Mecinas et al., 2011; Reul, 2014). In contrast, groups of genes that are regulated at 90 min are not only involved in transcriptional regulation but also in metabolic processes, such as maintenance of the internal state of glucose. This makes sense in light of the fact that the key purpose of the stress response is to supply the energy resources necessary to meet heightened metabolic demand (Floriou-Servou et al., 2021; Picard et al., 2018; Ulrich-Lai and Herman, 2009). Previous *in vitro* data had shown that application of the stress hormone corticosterone can trigger gene expression changes that last up to 5 hours. Our data show that all gene expression changes dissipate between 3-4 hours after stress, suggesting that regulation of transcription is very tightly controlled *in vivo*. This is in agreement with the notion that an efficient termination of the stress response is crucial for recovery (De Kloet et al., 2005; Floriou-Servou et al., 2021; Joëls and Baram, 2009).

### Stress effects are blunted on the level of translation

It is known that mRNA levels do not always correlate well with the levels of the corresponding proteins (Floriou-Servou et al., 2018; King and Gerber, 2014; Tian et al., 2004). In fact, evidence suggests that the control of protein abundance predominantly happens at the level of translation (Schwanhäusser et al., 2011). To address this, we employed TRAP-seq (Heiman et al., 2014), which affords cell type specificity, avoids cell dissociation, minimizes loss of RNAs that are localized in dendrites or axons, and allows a high depth of sequencing (Dougherty, 2017; King and Gerber, 2014). We extend previous work that has used TRAP-seq to reveal the translatomic response to AS specifically in excitatory neurons of the CA3 region (Gray et al., 2016), by providing the first global translatome profiling after stress, and a comparison of the translational profile for inhibitory and excitatory neurons of the dHC and vHC. In agreement with previous reports (Gutièrrez-Mecinas et al., 2011; Reul, 2014), *Egr1* and *Fos* were among the top upregulated targets, however *Egr1* was upregulated much more strongly in inhibitory neurons. Similarly, we found *Egr2* to be induced by stress mainly in inhibitory neurons, but in contrast *Cyr61* is strongly upregulated in excitatory neurons. In general, however, we detect far fewer stress-induced changes in the translatome, than expected based on our transcriptomic data. Importantly, our bona-fide stress-induced genes (Figure 6I) strongly agree with our recent meta analysis of published TRAP-seq data (von Ziegler et al., 2020), which gives us confidence that our analysis accurately captures the translatomic response.

It was particularly surprising that none of the translatomic changes we detected were observed on the level of the proteome. However, there are important technical limitations to consider. While our power analysis revealed a sensitivity similar to transcriptomics for quantifiable proteins, of all 1264 genes differentially altered on the transcriptional level, we were only able to quantify 105 corresponding proteins. This is a lower overlap than expected (expected observations = 294 proteins), and could be due to the fact that many of the significantly altered TFs are notoriously hard to detect using proteomics, owing to their low abundance and nuclear localization. However, when restricting the analysis to these proteins only, we found no increase in average significance either (data not shown). Subcellular fractionations might help to increase sensitivity and power, as some previous work suggests that long-lasting protein changes might be primarily restricted to synaptic compartments (Popoli et al., 2012). Despite these technical limitations, the absence of proteomic changes, together with a blunted translational response as assessed with TRAP-seq, is reminiscent of translational repression as it occurs in the context of the cellular integrated stress response (Advani and Ivanov, 2019; Pakos-Zebrucka et al., 2016). How the psychophysical stress response we describe here compares to the integrated stress response needs to be addressed in future work.

### Glial cell activation after acute stress

While the initial phospho-proteomic response seems to be dominated by neuronal/synaptic proteins, our data indicate that a sizable part of the molecular response can be attributed to glial cells. This supports a growing literature that astrocytes are key players in the stress response (Floriou-Servou et al., 2021; Murphy-Royal et al., 2019). Indeed, an elegant translatome screen in astrocytes from the somatosensory cortex revealed robust translational changes 90 min following acute restraint stress (Murphy-Royal et al., 2020). There is ample evidence that increased metabolic demand during stress requires rapid lactate shuttling from astrocytes to neurons (for review, see (Floriou-Servou et al., 2021)). In contrast, corticosterone signaling via glucocorticoid receptors, which are highly expressed in astrocytes, appears to limit astrocyte-to-neuron lactate shuttling (Murphy-Royal et al., 2020).

This explains how astrocytes become activated and inactivated in response to AS and also has important implications for chronic stress, which alters the interaction between astrocytes and neurons (Nasca et al., 2017). In addition, it explains why stress-related disorders are beginning to be considered as diseases of energy imbalance and metabolic dysregulation (Cherix et al., 2020; Picard and Sandi, 2021; Picard et al., 2018).

#### Summary and Outlook

Our work provides a detailed characterization of the molecular cascades that unfold as part of the healthy AS response. It also lays the foundation to further explore how stressors of different severity or duration impact different cell types and the metabolic profile of brain structures involved in mediating the stress response. To facilitate this process, all our data are publicly accessible and searchable through an interactive webportal https://bohaceklab.hest.ethz.ch/StressomeExplorer. Future studies will need to test whether more severely stressful experiences, or chronic stress exposures, lead to more exaggerated, less tightly regulated transcriptional and translational consequences, and whether such dysregulation would be associated with stress-related psychiatric disorders.

## Acknowledgements

The lab of JB is funded by the ETH Zurich, the ETH Project Grant ETH-20 19-1, SNSF Grant 310030_172889, Botnar Research Center for Child Health, 3R Competence Center, Kurt und Senta Herrmann-Stiftung, the Novartis Foundation for Medical-Biological Research and the Swiss Foundation for Excellence and Talent in Biomedical Research.

We thank the staff of the EPIC for the excellent animal care and their service to our animal facility. We thank Prof. Markus Stoffel for generously providing CMV-Cre mice. Parts of figures were created using Biorender.com.

**The authors declare no conflict of interest.**

## Abbreviations

S: Supplementary
AS: acute stress
HC: hippocampus
TF: transcription factor
min: minutes
IP: immunopurification

## Author contributions

**Lukas von Ziegler:** conceived experiments, conducted experiments & tissue processing (phosphoproteome immediate & time-course, time-course transcriptomics, proteome), data analysis, graphs & figures, interpreted results, wrote manuscript

**Amalia Floriou-Servou:** conceived experiments, conducted experiments & tissue processing (TRAP), graphs & figures, interpreted results, wrote manuscript

**Rebecca Waag:** assistance with TRAP sample processing, conducted single-nucleus RNA-seq

**Rebecca R. Das Gupta:** helped establish TRAP protocol, assistance in TRAP sample processing

**Oliver Sturman:** conducted experiments (behavior, assistance with time-course transcriptomics & immediate phosphoproteomics)

**Katharina Gapp:** conducted small-RNAseq

**Christina A. Maat:** produced western blots

**Tobias Kockmann:** assisted with the MS

**Han-Yu Lin:** assisted with sample processing

**Sian N. Duss:** conducted experiments (assistance with time-course phosphoproteomics, behavior)

**Mattia Privitera:** conducted experiments (behavior)

**Laura Hinte:** helped establish the snRNA-seq protocol; assistance with snRNA-seq

**Hanns U. Zeilhofer:** lab space and resources for 4 rounds of TRAP sample processing, advice

**Ferdinand von Meyenn:** lab space, expertise and resources for snRNAseq, advice

**Pierre-Luc Germain:** conducted data analysis and data integration, generated graphs & figures, interpreted results, wrote manuscript

**Johannes Bohacek:** conceived experiments, interpreted results, provided resources and funding, wrote manuscript

## Data availability

All LC-MS/MS data are available to readers via ProteomeXchange with identifier PXD024829. All sequencing data are available to readers via Gene Expression Omnibus with identifiers GSE169505 (bulk and TRAP sequencing), GSE169509 (smallRNA sequencing) and GSE169510 (snRNA sequencing). Code for the interactive web app and all analyses (independent scripts) can be found at https://github.com/ETHZ-INS/StressomeExplorer. Selected statistical results showing group contrasts and gene/protein clusters are reported in Table S1 (phosphoproteomics), Table S2 (transcriptomics) and Table S3 (proteomics).

## Methods and materials

### Animals

All experiments were conducted in accordance with the Swiss federal guidelines for the use of animals in research and under license ZH161/17, approved by the Zurich Cantonal veterinary office. For experiments with wild type animals, C57Bl/6J mice were either obtained from Janvier (France) or bred at the ETH Zurich animal facility (EPIC). For the generation of CMV-nuTRAP mice, heterozygous CMV-Cre mice (B6.C-Tg(CMV-cre)1Cgn/J, MGI:3613618) were bred with homozygous or heterozygous floxed nuTRAP mice (B6;129S6-Gt(ROSA)26Sor^tm2(CAG-NuTRAP)Evdr/J^, MGI:104735). For the generation of CaMKIIa-nuTRAP mice, homozygous floxed nuTRAP mice (B6;129S6-Gt(ROSA)26Sor^tm2(CAG-NuTRAP)Evdr/J^, MGI:104735) were bred with homozygous CaMKIIA-Cre (B6.Cg-Tg(Camk2a-cre)T29-1Stl/J, MGI:3613616) mice. The vGAT::bacTRAP (Tg(Slc32a1-RPL10a-eGFP)Uze) (Das Gupta et al., 2021) mice were generously provided by Prof. Hanns Ulrich Zeilhofer. Mice were housed in groups of 4-5 per cage in a temperature- and humidity-controlled facility on a 12-hour reversed light-dark cycle (lights off: 9:15 am; lights on: 9:15 pm), with food and water ad libitum, and used for experiments at the age of 2-5 months. All experiments were conducted during the animals’ active (dark) phase. Experiments were conducted with either male or female mice.

### Open field test (OFT)

Open-field testing was performed in sound insulated and ventilated multi-conditioning chambers (MultiConditioning System, TSE Systems Ltd, Germany), as described previously (Sturman et al., 2018). Briefly, the open field arena had dimensions 45 cm (l) x 45 cm (w) x 40 cm (h), and consisted of four transparent Plexiglas walls and a light gray PVC floor. Mice were tested under dim lighting (4 Lux) 75 dB of white noise was playing through the speakers of each box. Mice were placed directly into the center of the open field and the tracking/recording was initiated upon the first locomotion grid beam break. The test lasted 10 minutes.

### Elevated Plus Maze (EPM)

The EPM was made from gray PVC, with arms measuring 65.5 × 5.5 cm (*L* × *W*), elevated 61.5 cm.

Light intensity in the open arms was at 19–21 lux. All EPM tests were 5 minutes in duration. Distance, velocity, time in zone (open/closed arms + center) and head dips were recorded as described previously (Sturman et al., 2020).

### Swim stress paradigm and tissue collection

For all experiments, mice were single-housed 24 hours before exposure to stress. For cold swim stress, mice were placed for 6 min in a plastic beaker (20 cm diameter, 25 cm deep) filled with 18 ± 0.1°C water to 17 cm, in a room with dim red lighting. Immediately after stress exposure, mice returned to their assigned single-housing homecage. At the appropriate time point after initiation of stress, mice were euthanized by cervical dislocation and decapitation. The brain was quickly dissected on ice and isolated hippocampi were either snap-frozen in liquid nitrogen and stored at −80°C until further processing, or transferred to a binocular microscope and cooled with PBS-ice for dissection of subregions (proteomics) as previously described (von Ziegler et al., 2018) prior to snap-freezing and storing at −80°C.

### Unsupervised Clustering analyses for behavior

We generate temporally expanded pose-estimates as previously described (Sturman et al., 2020). All analyses were performed with RStudio (v.1.3.1073) using R (v. 3.6.3). The first and last 300 frames (=12 seconds) were removed to reduce noise from recording start and end. Temporally resolved pose estimates (± 15 frames) were first z-score normalized across all videos, then clustered using k-mean clustering with the bigkmeans() function from the “biganalytics” package (v1.1.21). Clustering was performed with 20 centers. Cross validation of clusters was performed with a random sub-set (5% of all sequences) of the same data with the tsne() function of the “M3C” package (v.1.8.0), plotting sequences by tsne 1 and tsne 2 coordinates and coloring by kmeans clusters. Clusters were smoothed over ± 5 frames and the number of on-offsets were determined for each individual animal. Clusters that were removed by smoothing or that did not occur in more than one animal were excluded. Example videos were generated by randomly picking 2 occurences of each cluster from each video where available. To assess the correlations of clusters the corrplot() function of the “corrplot” package (v0.84) was used with pearson’s correlation analysis. Statistical values for clusters were obtained with two sided student’s t-test.

### Proteomics

#### Protein extraction

We used a block design for sample processing. Samples were split into multiple blocks, containing one replicate of each condition and sub-region/region. Processing order within blocks was randomized, as was the order of blocks. Proteins were extracted from pooled hemispheres of dHC or vHC CA1, CA3 and DG samples using 150 μL TEAB buffer (100 mM triethylammonium bicarbonate, 0.1% SDS, 1:100 protease inhibitor cocktail P8340 (Sigma-Aldrich, St. Louis MO, USA), 1:500 PMSF (50 mM in EtOH)). Samples for phosphoproteomics (pooled hemispheres of vHC/dHC) were extracted using the same buffer with the addition of 1:100 phosphatase inhibitor cocktail 2 (P5726, Sigma-Aldrich, St. Louis MO, USA), and phosphatase inhibitor cocktail 3 (P0044, Sigma-Aldrich, St. Louis MO, USA).

The samples were mechanically lysed by 15 strokes with a 26 G needle and sonicated for 5 minutes. Samples were spun down at 16000 g for 30 minutes (1 hour for phospho samples) at 4°C and supernatants were collected. Proteins were quantified using a Qubit protein assay kit (ThermoFisher Scientific, Waltham MA, USA) following the manufacturer’s protocol. Protein extracts were further processed with a filter assisted sample preparation protocol (Wiśniewski et al., 2009). 20 μg of protein were added to 30 μL SDS denaturation buffer (4% SDS (w/v), 100 mM Tris/HCL pH 8.2, 0.1 M DTT). For phospho samples 300 μg of protein were added to equal amount (v/v) of concentrated denaturation buffer (0.2 M DTT). For denaturation, samples were incubated at 95°C for 5 minutes. Samples were diluted with 200 μL UA buffer (8M urea, 100 mM Tris/HCl pH 8.2) and then loaded to regenerated cellulose centrifugal filter units (Microcon 30, Merck Millipore, Billercia MA, USA). Samples were spun at 14000 g at 35°C for 20 minutes. Filter units were washed once with 200 μL of UA buffer followed by centrifugation at 14000 g at 35°C for 15 minutes. Cysteines were alkylated with 100 μL freshly prepared IAA solution (0.05 M iodoacetamide in UA buffer) for 1 minute at room temperature (RT) in a thermomixer at 600 rpm followed by centrifugation at 14000 g at 35°C for 10 minutes. Filter units were washed 3 times with 100 μL of UA buffer then twice with a 0.5 M NaCl solution in water (each washing was followed by centrifugation at 35°C and 14000 g for 10 minutes). Proteins were digested overnight at RT with a 1:50 ratio of sequencing grade modified trypsin (0.4 μg for whole proteome samples, 6 μg for phospho samples, V511A, Promega, Fitchburg WI) in 130 μL TEAB (0.05M Triethylammoniumbicarbonate in water). After protein digestion overnight at RT, peptide solutions were spun down at 14000 g at 35°C for 15 minutes and acidified with 10 μL of 5% TFA (trifluoroacetic acid). 100% ACN was added for a final concentration of 3% in the samples. Phospho-samples were split into reference samples (1:20 of final amount, used for peptidic clean-up) and enrichment samples (19:20 of final amount, used for phospho enrichment).

#### Enrichment of phospho peptides

For phospho enrichment digested samples were first lyophilized and resuspended in 200 μL loading buffer (1 M glycolic acid in 80% ACN, 5% TFA). Phosphopeptides were enriched using MagReSyn^®^ Ti-IMAC magnetic beads (Resyn Biosciences). 20 μL of beads were used per sample. Microspheres were washed once with 400 μL of 70% EtOH for 5 minutes at RT and reactivated using 200 μL of resuspension solution (1% NH4OH) for 10 minutes. Microspheres were then equilibrated twice with 200 μL loading buffer for 1 minute and then resuspended in 500ul 100% ACN per 20ul of beads. A Thermo Scientific™ KingFisher™ was used for enrichment of phospho peptides. First, microbeads were loaded onto magnetic rods and washed with 500 μL loading buffer. Samples were loaded and washed in 500 μL loading buffer, in 500 μL wash 2 buffer (80% ACN, 1% TFA) and in 500 μL wash 3 buffer (10% ACN, 0.2% TFA). Samples were then eluted in 200 μL resuspension solution (1% NH4OH) and lyophilized in a speedvac then re-solubilized in 19 μL 3% ACN / 0.1% FA (formic acid) prior to LC-MS/MS measurements. 1 μL of synthetic peptides (Biognosys AG, Switzerland) were added to each sample for retention time calibration.

#### Peptide clean-up

For non-phospho samples digests were cleaned-up using StageTip C18 silica columns (SP301, Thermo Scientific, Waltham MA). Columns were conditioned with 150 μL methanol followed by 150 μL of 60% ACN (acetonitrile) / 0.1% TFA. Columns were equilibrated with 2 x 150 μL of 3% ACN / 0,1% TFA. Samples were loaded onto the columns. They were then washed with 2 x 150 μL3% ACN / 0.1% TFA and eluted with 150 μL 60% ACN / 0.1% TFA. Samples were lyophilized in a speedvac then re-solubilized in 19 μL 3% ACN / 0.1% FA (formic acid) prior to LC-MS/MS measurement. 1 μL of synthetic peptides (Biognosys AG, Switzerland) were added to each sample for retention time calibration.

#### LC-MS/MS measurements

Samples were measured on a Q Exactive HF (Thermo Fisher Scientific, Waltham MA, USA). Peptides were separated with an ACQUITY UPLC M-Class System (Waters, Milford MA, USA). We used a single-pump trapping 75-μm scale configuration (Waters, Milford MA, USA). 1 μL of each sample (4 μL for phospho samples) were injected. Trapping was performed on a nanoEase^™^ symmetry C18 column (pore size 100Å, particle size 5 μm, inner diameter 180 μm, length 20 mm). For separation a nanoEase^™^ HSS C18 T3 column was used (pore size 100Å, particle size 1.8 μm, inner diameter 75 μm, length 250 mm, heated to 50°C). Peptides were separated using a 120 minute long linear solvent gradient of 5-35% ACN, 0.1% FA or a 60 minutes long linear gradient for phospho peptides (both using a flowrate of 300 nL / minute). Electronspray ionization with 2.6 kV was used and a DIA method with a MS1 in each cycle followed by 35 fixed 20 Da precursor isolation windows within a precursor range of 400-1100 m/z was applied. For MS1 we used a maximum injection time of 55 ms and an AGC target of 3e6 with a resolution of 30 K in the range of 350-1500 m/z. MS2 spectra were acquired using a maximum injection time of 55 ms an AGC target of 1e6 with a 30 K resolution in the range of 140 – (2 x the upper range of the precursor window) m/z. A HCD collision energy of 28 was used for fragmentation.

#### Peak picking and quantification of non-phospho samples

We used Spectronaut^™^ (Biognosys, version 10) with directDIA for peak picking and sequence assignment. We used a *Mus musculus* reference proteome for C57BL/6J from uniprot (UP000000589) from the Ensembl GCA_000001635.8 assembly only including reviewed entries. We included a maximum of 2 missed cleavages, using a Tryptic specificity (KR/P). Sequences in a range of 7-52 AA were considered. We included carbamidomethyl as fixed modification for cysteine, oxidation as variable modification for methionine and protein N-terminal acetylation as variable modification. A maximum of 5 variable modifications were considered. Single hit was determined on the stripped sequence level. Major grouping was done by protein group ID and minor grouping by stripped sequence. Only proteotypic peptide sequences were considered and single hit proteins excluded. For the minor and major group quantification the top 3 entries were used using the mean precursor/peptide quantity. A localized normalization strategy and interference correction were used. Machine learning was performed on a per run basis and iRT profiling was enabled.

#### Peak picking and quantification of phospho samples

For phospho samples we used following additions/modifications to the Spectronaut^™^ searches. We further included phosphorylation of serine, threonine or tyrosine as variable modification, either as full group or following a neutral loss of 98. PTM localization was enabled with a probability cutoff of 0.75 and quantification performed on the precursor level. We included both proteotypic and non-proteotypic peptides. A global median normalization strategy was used.

#### Statistical analysis of LC-MS/MS data

For non-phospho samples we used exported quantity values of proteins from Spectronaut. For statistical analysis we used R version 3.6.2. We performed 2 group analyses (Homecage vs Swim) within subregions and regions. We used log2 transformed quantity values and performed an ANOVA analysis (using the aov() of function R) whilst correcting for block effects. PCA was performed on the expression matrix of all samples. For subregion vs. region analysis we used a two-way ANOVA with the formula sub-region * region while correcting for overall effects within animals.

For phospho measurements quantity values were exported from Spectronaut on the modified peptide level. We then used the R package “DEP” (version 1.6.1) for differential analysis of defined contrasts. For the missing value filter we used a threshold of 2 and for the imputation we used the “MinProb” method with a q = 0.01.

For multiple testing correction the Benjamini-Hochberg false discovery rate (FDR) method was used.

### Transcriptomics and translatomics

#### Whole tissue RNA extraction

We used a block design for sample processing. Samples were split into multiple blocks, containing one replicate of each condition and region and processing order within blocks randomized. Samples were homogenized in 500 μL Trizol (Invitrogen 15596026) in a tissue lyser bead mill (Qiagen, Germany) at 4°C for 2 minutes, and RNA was extracted according to manufacturer’s recommendations. RNA purity and quantity were determined with a UV/V spectrophotometer (Nanodrop 1000), while RNA integrity was assessed with high sensitivity RNA screen tape on an Agilent Tape Station/Bioanalyzer, according to the manufacturer’s protocol. The RIN values of all samples ranged from 8.4 to 10.0. For library preparation, the TruSeq stranded RNA kit (Illumina Inc.) was used according to the manufacturer’s protocol.

#### Processing of TRAP samples

To isolate actively translated RNA from specific cell types in the HC, the TRAP protocol was followed as described in (Heiman et al., 2014). The samples were homogenized with a 2 mL tissue grinder and a pestle (3432S90, Thomas Scientific), on a Rotor for homogenizers (LT-400D, Yamato) in tissue lysis buffer. Solutions and the affinity matrix were prepared as recommended, and optimal bead titers were determined for every transgenic line with affinity matrix titration. More specifically, for samples of both the dHC and vHC of CaMKIIa-nuTRAP mice, the affinity matrix was prepared using 75 μL of Streptavidin MyOne T1 Dynabeads (65601, Invitrogen) per sample. For the vGAT::bacTRAP mice 75 μL beads per sample were used for the dHC and 150 μL for the vHC, and for the vHC samples of the CMV-nuTRAP mice 300 μL beads were used per sample. The corresponding volumes of biotinylated protein L and the GFP antibodies 19C8 and 19F7 were added, and the affinity matrix (in a final volume of 200 μL) was added to every sample for immunopurification. Samples were incubated at a tube rotator at 4°C for roughly 17 hours.

After immunopurification we proceeded to RNA elution and purification with the RNeasy Plus Micro Kit (74034, Qiagen). The RNA was first dissociated from the ribosomes/beads with 350 μl Buffer RLT Plus from the kit, containing 40 μM dithiothreitol (DTT, R0861, Thermo Fisher Scientific), and purified according to the manufacturer’s instructions. RNA concentration was estimated with either the RiboGreen fluorescence assay (R11491, Thermo Fisher Scientific) and a NanoDrop 3300 microvolume fluorospectrometer, or with the Quantifluor RNA System (E3310, Promega) and a DS-11 Series Spectrophotometer / Fluorometer (DeNovix). The integrity of the RNA was assessed using the RNA 6000 Pico Kit (5067-1513, Agilent Technologies) on a 2100 Bioanalyzer (Agilent Technologies Inc., CA), and all RIN values were between 8.2 and 10.0. Conversion to cDNA and generation of libraries were performed with the Smart-seq2 protocol (Picelli et al., 2014).

#### Bulk and TRAP sequencing and data analysis

Library preparation and sequencing was performed at the Functional Genomics Center Zurich (FGCZ) core facility. For bulk sequencing library preparation, the TruSeq stranded RNA kit (Illumina Inc.) was used according to the manufacturer’s protocol. The mRNA was purified by polyA selection, chemically fragmented and transcribed into cDNA before adapter ligation. For bulk and TRAP sequencing single-end sequencing (100nt) was performed with Illumina Novaseq or HiSeq 4000. Samples within experiments were each run on one or multiple lanes and demultiplexed. A sequencing depth of ~20M reads per sample was used. Adapters were trimmed using cutadapt (Martin, 2011) with a maximum error rate of 0.05 and a minimum length of 15. Kallisto (Bray et al., 2016) was used for pseudoalignment of reads on the transcriptome level using the genecode.vM17 assembly with 30 bootstrap samples and an estimated fragment length of 200 ± 20. For differential gene expression (DGE) analysis we aggregated reads of protein coding transcripts and used R (v. 3.6.2) with the package “edgeR” (v 3.26.8) for analysis. A filter was used to remove genes with low expression prior to DGE analysis. EdgeR was then used to calculate the normalization factors (TMM method) and estimate the dispersion (by weighted likelihood empirical Bayes). For two group comparisons the genewise exact test was used, for more complex designs we used a generalized linear model (GLM) with empirical Bayes quasi-likelihood F-tests. For multiple testing correction the Benjamini-Hochberg false discovery rate (FDR) method was used.

To quantify spliced (i.e. exonic) and unspliced transcripts, we aligned the reads using STAR (Dobin et al., 2013) and ran featureCounts (Liao et al., 2014) with three sets of parameters: i) using exons as features (standard), ii) using exons with --nonSplitOnly, and iii) using transcripts as features with --nonSplitOnly (all three additionally shared the “-O --largestOverlap --nonOverlap 3 --fracOverlap 0.9 --primary -s 2” parameters). The first was used as a quantification of processed transcripts; for unprocessed transcripts, we subtracted from the third the counts of the second (which are compatible with processed transcripts as well).

For analyses of data-sets originating from multiple experiments (i.e. different TRAP protocols) we further employed SVA correction to correct for processing specific effects (Leek et al., 2012). Surrogate variables independent of experimental groups were identified using the sva package 3.34.0 on data after DESeq2 variance-stabilization (Love et al., 2014), and were then included as additive terms in the GLMs.

For the metaanalysis of TRAP data across datasets (Figure 6I/J) we used the following SVA correction models and subsequent GLM models. additive analysis; SVA model: ~ *Dataset + Condition*, GLM model ~*SVs + Condition*. Interactive analysis; SVA model: ~ *Dataset * Condition*, GLM model: ~*SVs + Condition * Dataset*. SVs = surrogate variables

Heatmaps were produced with the SEtools package. To avoid rare extreme values from driving the scale, the color scale is linear for values within a 98% interval, and ordinal for values outside it.

#### Single nucleus RNA sequencing and analysis

Tissue dissociation: All steps for tissue dissociation were performed on ice. Both hippocampi of the same animal were chopped briefly in 100 μL lysis buffer (10 mM Tris-HCl, 10 mM NaCl, 3 mM MgCl2, 0.1% Igepal, 0.2 U/ul recombinant RNase inhibitor in nuclease-free water) and immediately transferred to a dounce homogenizer filled with 1 mL lysis buffer. Tissue was homogenized by 15-20 strokes with each pestle A and pestle B. 2 mL of lysis buffer was added to the dounce homogenizer and the solution was incubated for 5 minutes. The dounce homogenizer was filled with 4 ml wash buffer (1% BSA. 0.2 U/ul recombinant RNase inhibitor in PBS) and homogenate was passed through a 40 μm filter before adding 8 ml wash buffer. The sample was centrifuged at 500 g for 5 minutes at 4 °C and the pellet was resuspended in 7 mL nuclei suspension buffer (1% BSA, 0.2 U/μl RNaseInhibitor (Superasin) in PBS). Then the sample was centrifuged at 500 g for 5 minutes at 4 °C. The pellet was resuspended in 1 mL nuclei suspension buffer and filtered through a 40 μm, 30 μm and 10 μm filter. Nuclei count was assessed manually by Trypan blue staining and adjusted to a final concentration of 1000 nuclei/μL.

Library preparation and sequencing: cDNA synthesis and library preparation were conducted following the Chromium Single Cell 3’ Reagent Kit v3 User Guide (CG000183 Rev B, 10X Genomics). 14 PCR cycles were performed for cDNA amplification. Each library was sequenced on a NovaSeq6000 instrument aiming for a sequencing depth of 40’000 reads/nucleus.

For data analysis, reads were demultiplexed, aligned and quantified with Cell Ranger 4.0.0 (Zheng et al., 2017) using the GRCm38/GENCODE M23 pre-mRNA transcriptome from 10x. The code for downstream analysis is available in the repository. In brief, we first decontaminated the cells from ambient RNA using *SoupX* 1.4.5 (Young and Behjati, 2020), using cell clusters obtained from the “fastcluster” function of *scDblFinder 1.4.0*. We then removed doublet identified by *scDblFinder* (Germain et al., 2020), and additionally removed cells (using *scater 1.18.0* (McCarthy et al., 2017)) which had more than 9% mitochondrial reads or departed from the median by more than 3 median absolute deviations on either of the following cell QC metrics: log(UMI counts), log(features detected), or percent of UMI in the top 50 features (filtering out higher only). We next normalized the data using *sctransform 0.3.1* (Hafemeister and Satija, 2019), and ran PCA and clustering at various resolutions using *Seurat 3.2.2* (Stuart et al., 2019). For each resolution, we then trained an *xgboost* classifier (10-fold cross-validation at 7:3 train:test split) using the top 1000 genes to estimate the clusters’ stability and identifiability, identifying a Leiden resolution parameter of 0.25 as providing an appropriate resolution with low misclassification rate.

The clusters showed very distinct profiles for known markers of broad cell classes. To refine the annotation, we applied cell-level assignment using *scClassify 1.2.0* (Lin et al., 2020) (ensemble approach, using limma, BI and DD for feature selection, and Pearson, Spearman and Cosine as similarity metrics) based on the HCL1 reference from (Zeisel et al., 2018). There was overall a good agreement between our clusters and the cell-level assignments (in particular separating dentate gyrus neurons from other excitatory neurons), except for subtypes of vascular cells (which were represented in small numbers in our data). For this reason, for downstream analysis we kept this resolution for these cells. One cluster had cells with very low library sizes and showed expression of both neuronal and astrocytic markers, and was consequently excluded; in addition, two clusters with very low abundance had conflicting marker expression and were not represented across all samples; they were also excluded.

As preliminary differential expression analyses did not show significant differences in the response of the two astrocyte clusters or of the three excitatory neuron clusters, they were respectively merged as Astrocytes and Excitatory Neurons. Differential expression analysis was then performed using *muscat* 1.4.0 (Crowell et al., 2020) pseudo-bulk method. To assign bulk DEGs to a cell type on the basis of the single-nuclei data, we first calculated the absolute (pseudo-bulk) difference between stressed and control animals for each cell type, weighted it by multiplying it by the power of 1 minus the p-value in the cell type, and identified the cell type with the highest weighted difference in the same direction as the bulk change. When the absolute weighted difference in the top cell type was less than twice as high as the next cell type, the gene was assigned was considered ambiguous.

#### Transcription factor activity inference

Transcription factor regulons were obtained from (Garcia-Alonso et al., 2019), lifted to the mouse genome using ensembl orthologs, and the likelihood of the interactions was weighted according to the approximative AUC of the different categories in their benchmark. Category E interactions were discarded, and transcription factors that did not pass differential expression filtering were excluded. We next calculated sample-wise activity score per transcription factor in the unspliced data using *viper* 1.24 with pleiotropy correction (Alvarez et al., 2016). To identify factors with differences in activity, we then fitted ~region+timepoint models on each factor and tested for dropping the time point coefficients using *limma* 3.46 (Ritchie et al., 2015). Only factors with an FDR < 0.005, an absolute log-fold change > 0.2 and a mean log(counts per million) >= 2 were considered.

To co-cluster differentially-expressed genes based on both regulons and expression, we first produced a TF-target matrix multiplying the likelihood of the interaction with the factor’s mode of action, and performing z-score normalization on each factor, and merging this matrix with the genes’ log-fold changes at 45 minutes, 90 minutes and 2 hours, in both unspliced and spliced fractions. Partitioning around 9 medoids was used to cluster genes based on this merged information. For each cluster, gene ontology enrichment analysis was performed with *goseq* 1.41 (Young et al., 2010) using the tested genes as background, and the top terms were reported after eliminating entirely redundant terms. Overlayed protein-protein interactions between the selected factors were obtained from the BioGrid 4.2.193 (Oughtred et al., 2019).

#### Small RNA sequencing and data analysis

1000 ng of total RNA was used as input for Nextflex v3 small RNA library preparation (Perkin Elmer Applied Genomics). Adapters and primers were undiluted and libraries were amplified with 15 PCR cycles prior bead-based size selection. Libraries were prepared in 3 batches composed of samples of both hippocampal subregions and stressed/non-stressed groups. All resulting libraries were pooled to be equimolar and sequenced on 2 lanes of an Illumina Genome Analyzer HiSeq 2500 (Illumina) with a single ended read length of small 50 bp. The experimenter was blinded to the treatment.

For data analysis, reads were first trimmed using cutadapt (adapter sequence TGGAATTCTCGGGTGCCAAGG, 0.05 error rate, minimum length of 18). Reads with the same sequence (including UMI) were then collapsed, and the UMIs trimmed. Quantification was then performed using Sports 1.0 (Shi et al., 2018), using the software’s full mouse annotation as described. SVA analysis was used to identify two vectors of technical variation which were then included, along with the hippocampal region, as covariates in the differential expression model. Differential expressions testing was done with edgeR using QLFit, testing for the dropping of the condition and region:condition coefficients.

#### Tissue processing and immunohistochemistry

In order to confirm the localization of fluorescent tags in excitatory neurons in the CaMKIIa-nuTRAp mice, in inhibitory neurons in the vGAT::bacTRAP mice, and in both neurons and glia in CMV-nuTRAP mice we performed immunohistochemistry as previously described (Zerbi et al., 2019). More specifically, the mice were first deeply anesthetized with pentobarbital (150 mg/kg, i.p.) and perfused intracardially through the left ventricle for 2 minutes, with approximately 20 mL of artificial cerebrospinal fluid (ACSF) or ice-cold PBS pH 7.4. Then the brain was dissected, blocked and fixed for 1.5-3 hr in ice-cold paraformaldehyde solution (4% PFA in PBS, pH 7.4). The tissue was cryo-protected, frozen in tissue mounting medium (Tissue-Tek O.C.T Compound, Sakura Finetek Europe B.V., Netherlands), and sectioned coronally into 40 μm thick sections.

The brain sections were incubated in primary antibody solution containing 0.2% Triton X-100, and 2% normal goat serum in PBS, at 4°C under continuous agitation, overnight. After 3 washes with PBS the sections were transferred in secondary antibody solution with 2% normal goat serum in PBS. Then the sections were washed again in PBS, mounted onto glass slides (Menzel-Gläser SUPERFROST PLUS, Thermo Scientific), air-dried and coverslipped with Dako fluorescence mounting medium (Agilent Technologies). For anti-GAD65/67, an additional step of antigen retrieval and one blocking step preceded the primary antibody incubation. During antigen retrieval, the sections were submerged in 500 μL/well citrate buffer 10 mM, pH 6, containing 0.05% Tween20 and remained there for 20 minutes, under soft agitation (30 rpm). Then the sections were rinsed with PBS and incubated in 500 μL/well blocking solution (5% NGS, 0.3% TritonX-100 in PBS), under soft agitation, at RT for 60 minutes. The primary antibody solution contained 0.3% Triton X-100, and 5% normal goat serum, and the secondary antibody solution contained 0.05% Triton X-100, and 5% normal goat serum in PBS.

Primary antibodies used: chicken anti-GFP (ab13970, Abcam, 1:1000), chicken anti-GFP (GFP-1020, Aves, 1:5000), rabbit anti-mCherry (ab167453, Abcam, 1:1000), rabbit anti-GAD65/67 (AB1511, Millipore, 1:500), rabbit anti-NeuN (EPR12763, ab177487, Abcam, 1:300), mouse anti-GFAP (G3893, Sigma-Aldrich, 1:400). Secondary antibodies used: goat anti-chicken Alexa Fluor 488 (A-11039, Thermo Fischer Scientific, 1:1000), goat anti-rabbit Alexa 546 (A11035, Thermo Fisher Scientific, 1:300), goat anti-mouse Cy™3 (115-165-003, Jackson ImmunoResearch, 1:300), goat anti-rabbit Alexa Fluor Plus 647 (A32733, Invitrogen, 1:500). Stains used: Nissl stain (N21483, NeuroTrace 640/660 Nissl stain, Thermo Fischer Scientific, 1:300), DAPI (D3571, Thermo Fischer Scientific, 1:2000). Microscopy images were acquired in a confocal laser-scanning microscope (CLSM 880, Carl Zeiss AG, Germany), maintaining a pinhole aperture of 1.0 Airy Unit and image size 1024×1024 pixels. Images were acquired using a Z stack with a 20x objective and pixel size 0.59 mm.

### Western blot

Tissue was mechanically homogenized in 200 μL TEAB lysis buffer and sonicated for 2 minutes at maximum intensity. Following centrifugation at 16000 g at 4°C for 30 minutes, the recovered supernatant was collected and the protein concentration was determined using the Qubit Protein Assay Kit (Q33211, Thermo Fischer Scientific). Proteins were denatured by boiling at 95°C in SDS-loading buffer containing 10% mercaptoethanol, for 5 minutes. 20 μg of total protein/sample were loaded in each well of a 10% PAGE gel and transferred onto a nitrocellulose membrane (#1704158, BioRad) in a Trans-Blot Turbo system (Bio-Rad). Membranes were blocked in a 5% non-fat milk solution in 0.1% Tween20 (TBS-T), for 1 hour at RT and incubated in primary antibody solution overnight at 4°C, or for 2 hours at RT. Subsequently, membranes were washed 3 times in TBS-T, incubated in secondary antibody solution at RT for 1 hour, washed again 3 times in TBS-T and visualized in a ChemiDoc Imaging System (Bio-Rad). Antibodies were diluted in antibody solution containing 1% non-fat milk in TBS-T. Primary antibodies used are the anti-pS553-Syn1 rabbit monoclonal (ab32532, Abcam, 1:10000) and anti-GAPDH rabbit polyclonal (ABS16, Millipore, 1:1000). The primary antibodies were targeted with the secondary antibody goat anti-rabbit IRDye800CW (926-32211, LI-COR Biosciences GmbH, 1:10000).

### RT-PCR

Real-Time polymerase chain reaction (RT-PCR) was performed using SYBR green (04887352001, Roche) on a CFX384 Touch Real-Time PCR Detection System (Bio-Rad). As described before (Roszkowski et al., 2016), the cycling protocol was: 5 min at 95°C, then 45 cycles each including a step of denaturation (10 s at 95°C), a step of annealing (10 s at 60°C), and a step of elongation (10 s at 72°C). Primers were designed with Quantprime (Arvidsson et al., 2008) and tested for quality and specificity by melt-curve analysis and gel electrophoresis, unless stated otherwise. The primers for AtLAS were designed by (Ma et al., 2020).

Forward (FP) and reverse (RP) primer sequences:

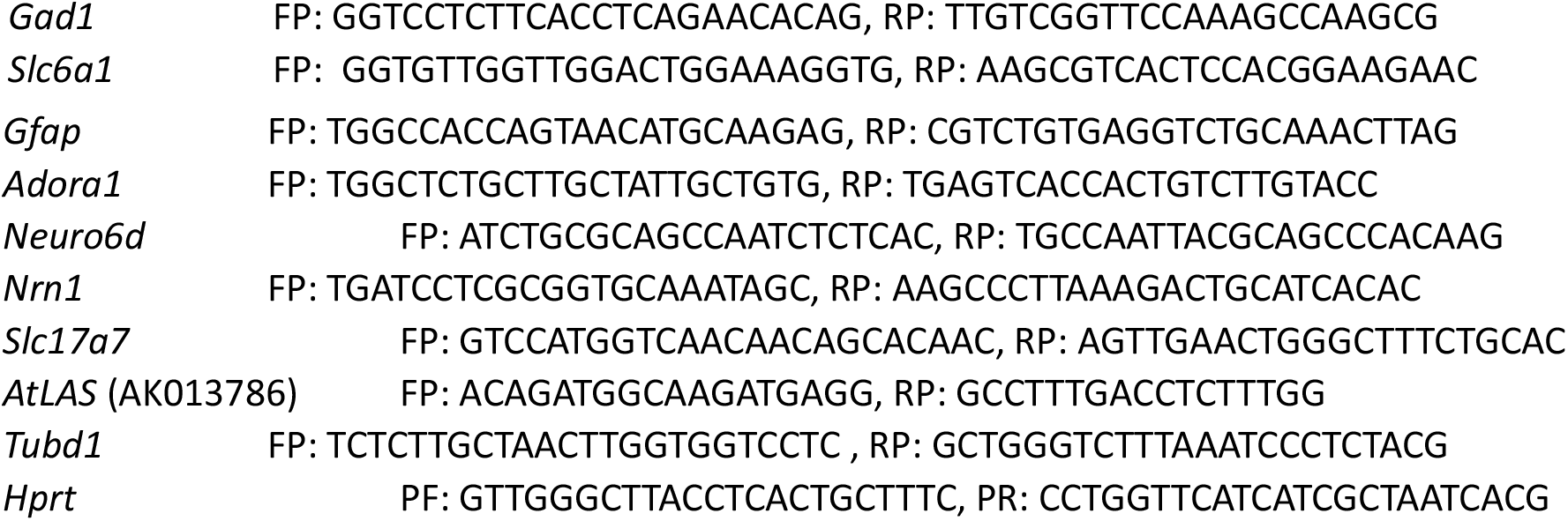

## Supplementary Figures

**Figure S1:**
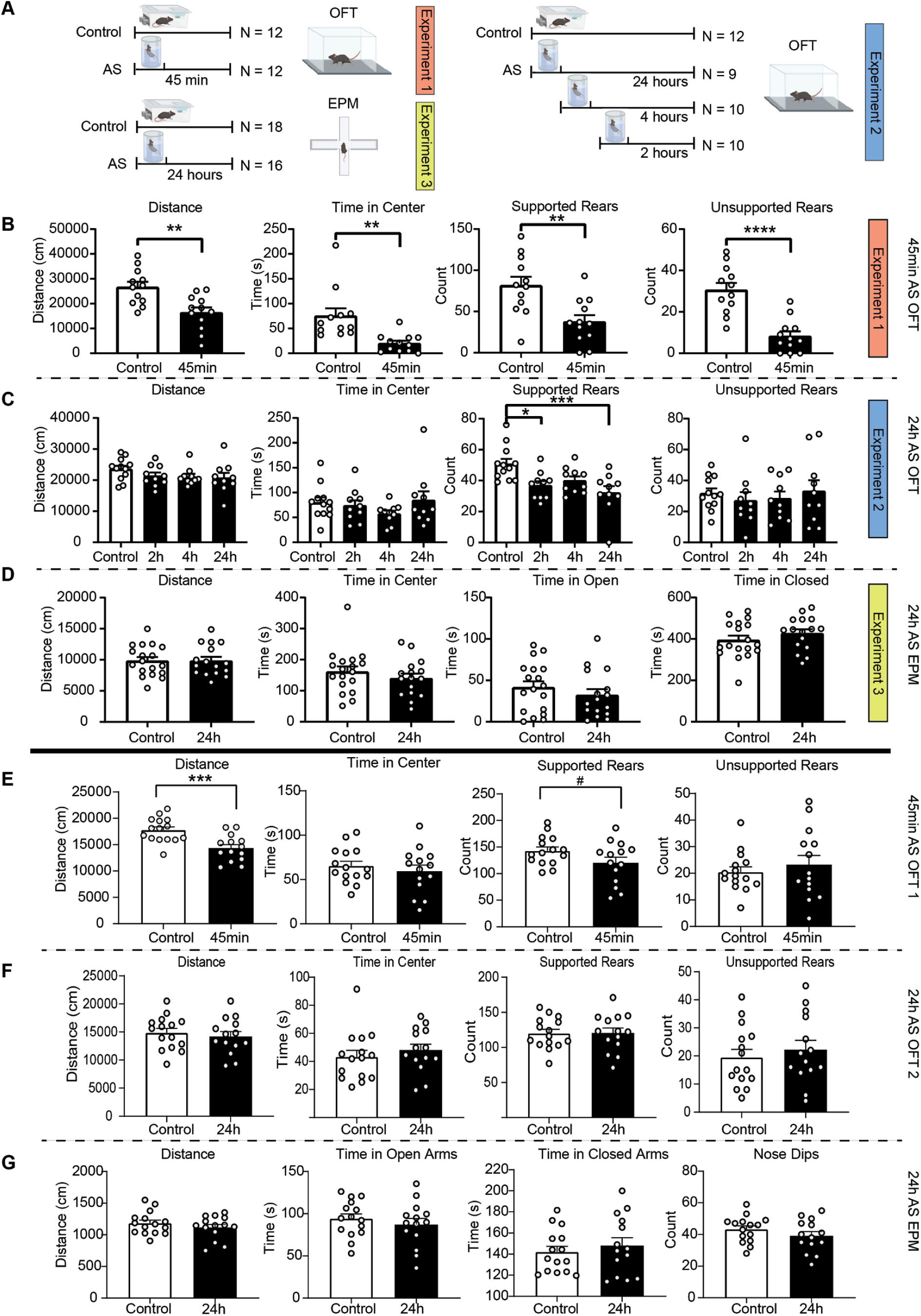
Stress induced changes in behavior across multiple experiments: **A)** Overview of 3 different experiments to assess stress induced behavioral changes at different time-points. **B)** Stress induced changes in the OFT after 45 minutes. Stressed animals show reduced distance traveled (unpaired t-test; t=3.548, df=22, p=0.0018), reduced time in center (unpaired t-test; t=3.5, df=22, p=0.0020) as well as reduced supported (unpaired t-test; t=3.393, df=22, p=0.0026) and unsupported (unpaired t-test; t=5.532, df=22, p<0.0001) rearing compared to controls (Ctrl N = 12; 45min swim N = 12). **C)** Stress induced changes in the OFT after 2, 4 and 24h. Stressed animals show reduced supported rearing after 2h and 24h (One-way ANOVA; F(3,38)=6.122, P=0.0017. Dunnett’s multiple comparisons test; Control vs 2h: adjusted p value 0.0115, Control vs 4h: adjusted p value=0.0007), however no change in distance traveled, time in center and unsupported rearing (Ctrl N = 12, 2h swim N = 10, 4h swim N = 10, 24h swim N = 24) **D)** Stress induced changes in the EPM after 24h. There are no differences between control and swim animals in distance traveled, time in center, time in open arm and time in closed arm. (Ctrl N = 18, 24h swim N = 18) **E)** stress induced classical readouts in the OFT after 45min in the unsupervised clustering experiment (Figure 1A). Stressed animals show reduced distance traveled (unpaired t-test; t=3.859, df=27, p=0.0006, and a trend of reduced supported rearing (unpaired t-test; t=1.789, df=27, p=0.0849)(control N = 15; swim N = 14). **F)** stress induced classical readouts in the OFT after 24h in the unsupervised clustering experiment (Figure 1A). No significant differences can be observed between control and stressed animals (control N = 15; swim N = 14). **G)** stress induced classical readouts in the EPM after 24h in the unsupervised clustering experiment (Figure 1A). No significant differences can be observed between control and stressed animals (control N = 15; swim N = 14). # p< 0.1, *p < 0.05, ** p < 0.01, *** p < 0.001, **** p < 0.0001, bars depict mean ± SEM.

**Figure S2:**
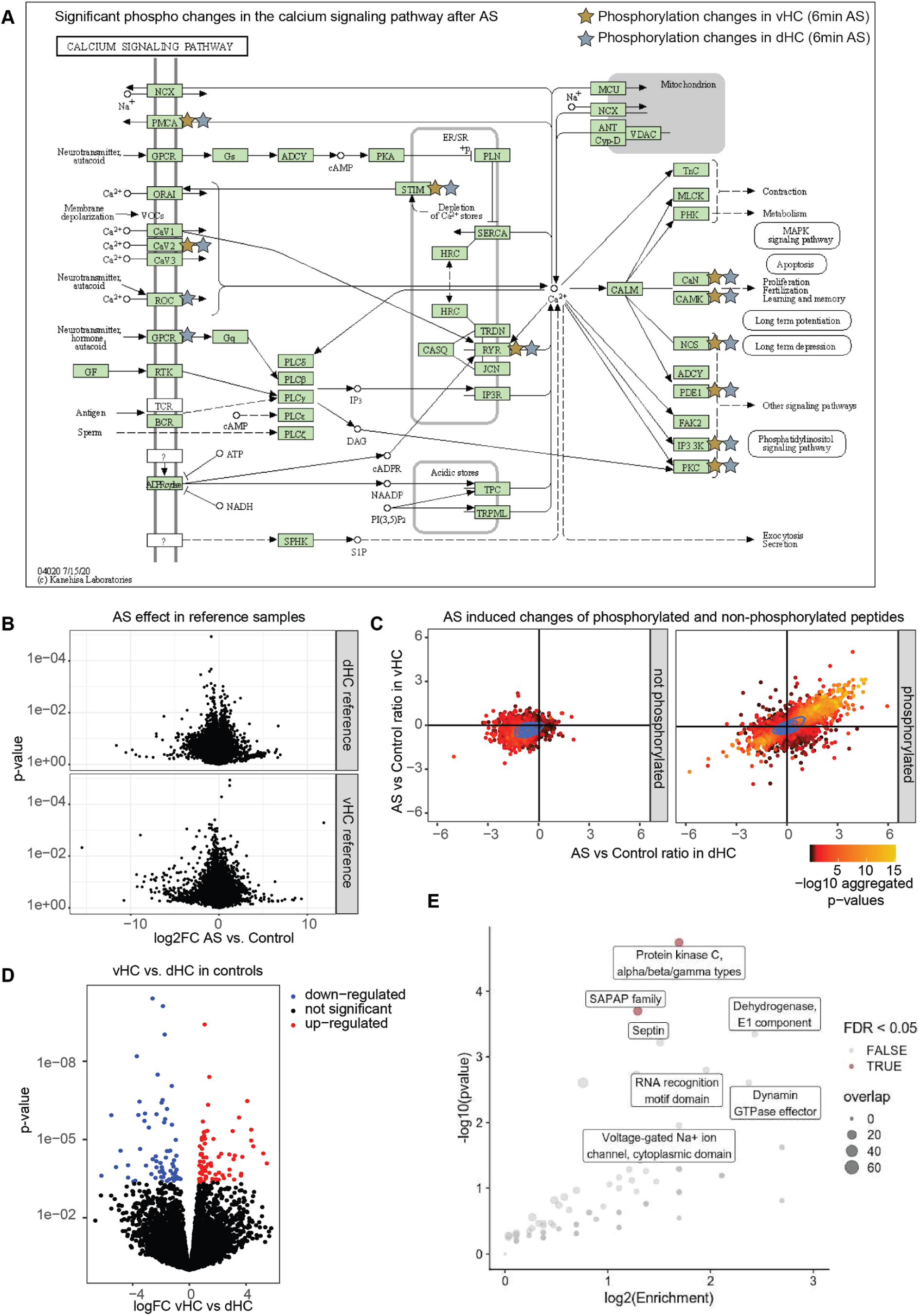
The phosphoproteome of acute swim stress. **A)** Calcium signaling pathway proteins with altered phosphorylation immediately after stress in vHC and dHC. Red stars label proteins which have upregulated phosphosites after acute stress. Original illustration adapted from kegg pathways (www.genome.jp/kegg) **B)** Both in dHC and vHC no significant changes are observed in reference samples (N = 4 × 4) after multiple testing correction, indicating no stress induced change of total protein expression. **C)** Strength of swim effects (6 minutes time-point) on phosphorylated vs non-phosphorylated peptides in vHC and dHC. **D)** Volcano plots depicting vHC vs dHC baseline differences in the control group **E)** Over-representation of InterPro protein domains among significantly altered phosphopeptides. Significantly overrepresented domains (FDR < 0.05) are labeled in red.

**Figure S3:**
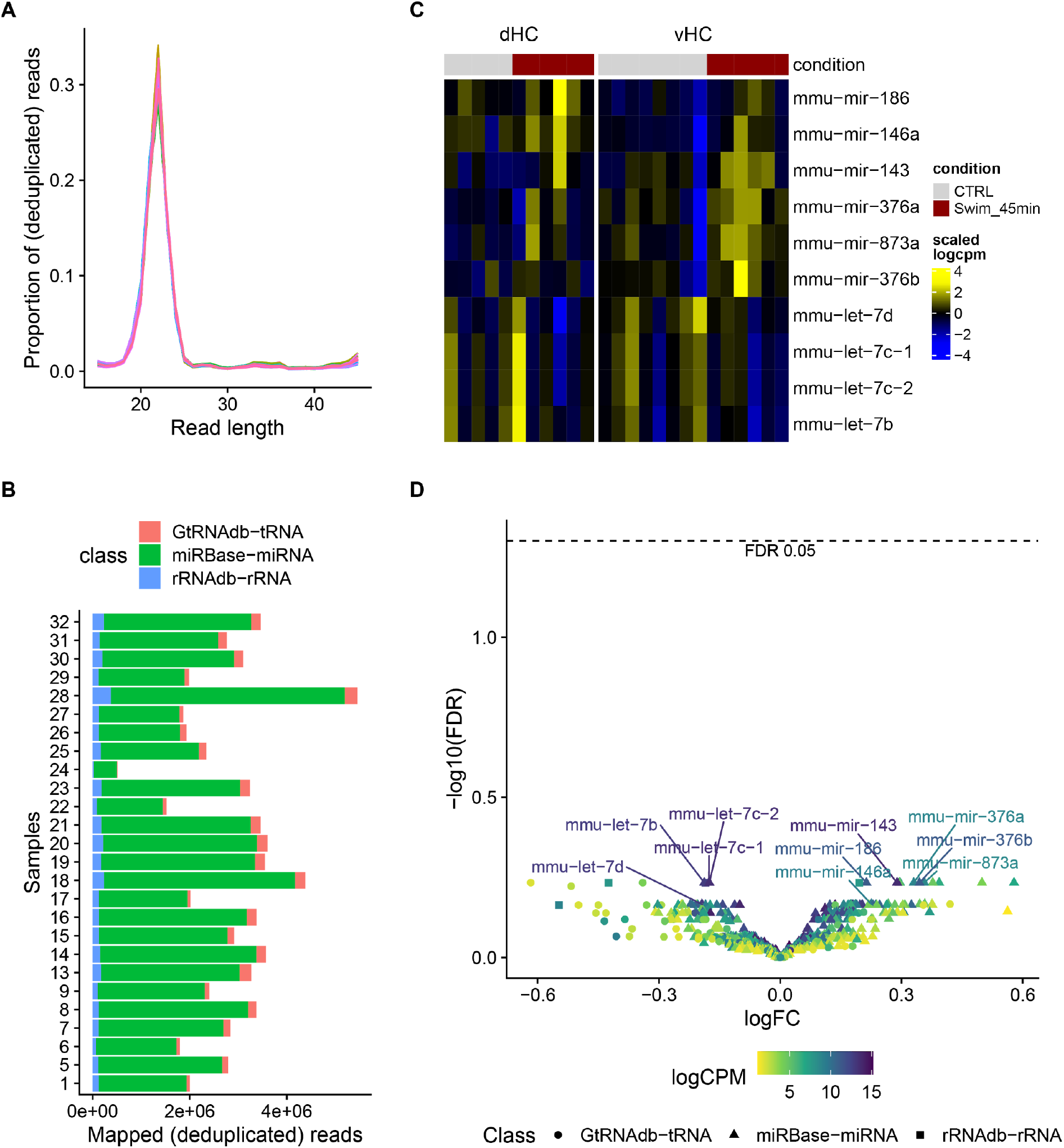
miRNA sequencing. **A)** Read length distribution across samples after UMI and adapter trimming. **B)** Mapped (deduplicated) reads assigned to the major small RNA classes. Sample 24 was excluded from downstream analysis due to low coverage. **C)** Relative expression profile of the top candidate miRNAs (not passing genome-wide significance). **D)** Volcano plot of the differential expression results.

**Figure S4:**
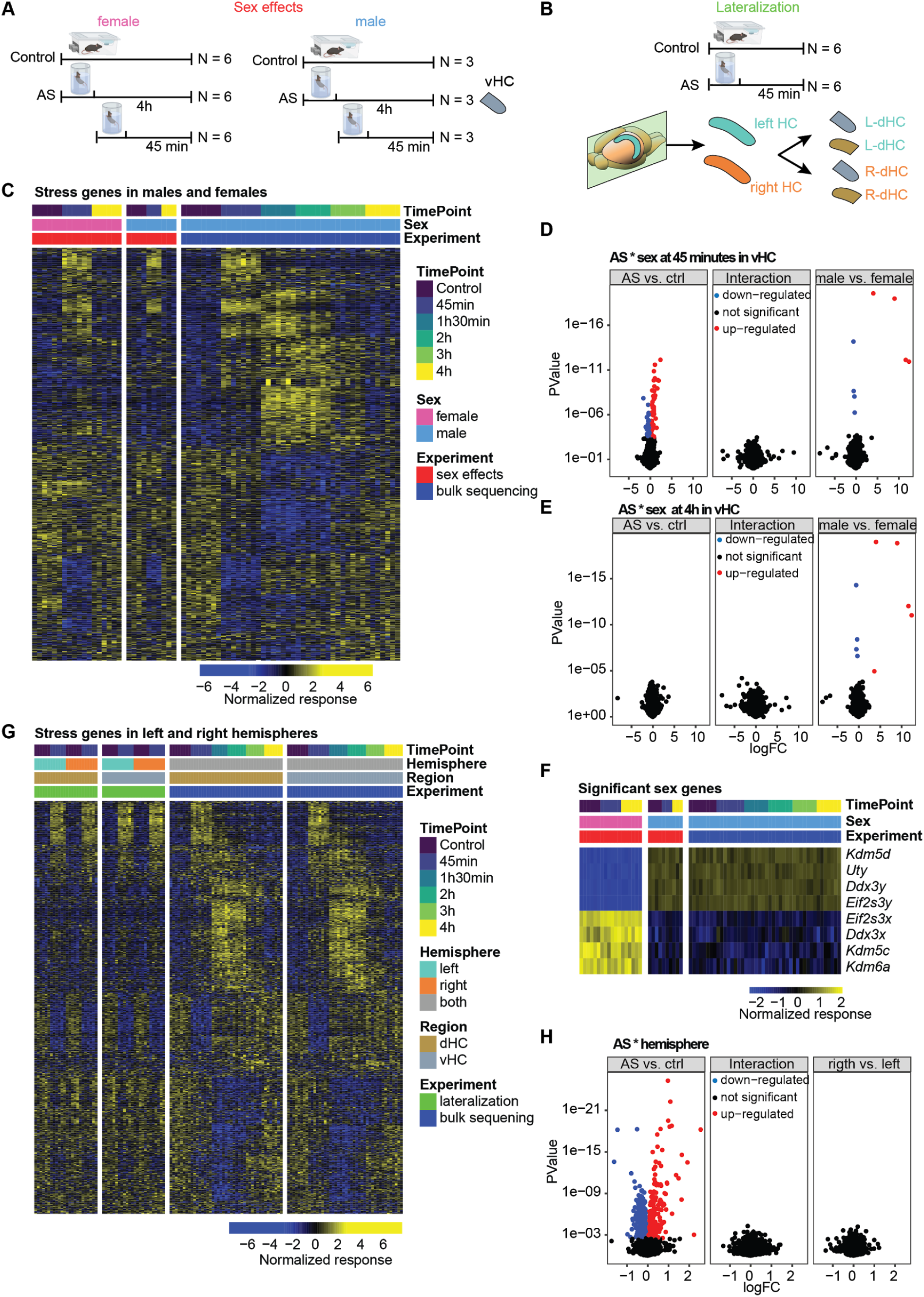
Transcriptomics effects of swim stress on the hippocampus of males and females and in the left and right hemispheres. **A)** Experimental design of experiments testing for sex differences. **B)** Experimental design of the lateralization experiment. **C)** Stress responsive genes from the bulk sequencing experiment in male and female vHC at 45 minutes and 4 hours. **D)** Volcanoplots depicting statistical results of stress and sex effects and sex:stress interactions at 45 minutes in the vHC. **E)** Volcanoplots depicting statistical results of stress and sex effects and sex:stress interactions at 4 hoursin the vHC. **F)** Heatmap of significant sex genes. **G)** Stress responsive genes from the bulk sequencing experiment left and right hemispheres. **H)** Volcano plots depicting statistical results of stress and hemisphere effects and hemisphere:stress interactions at 45 minutes.

**Figure S5:**
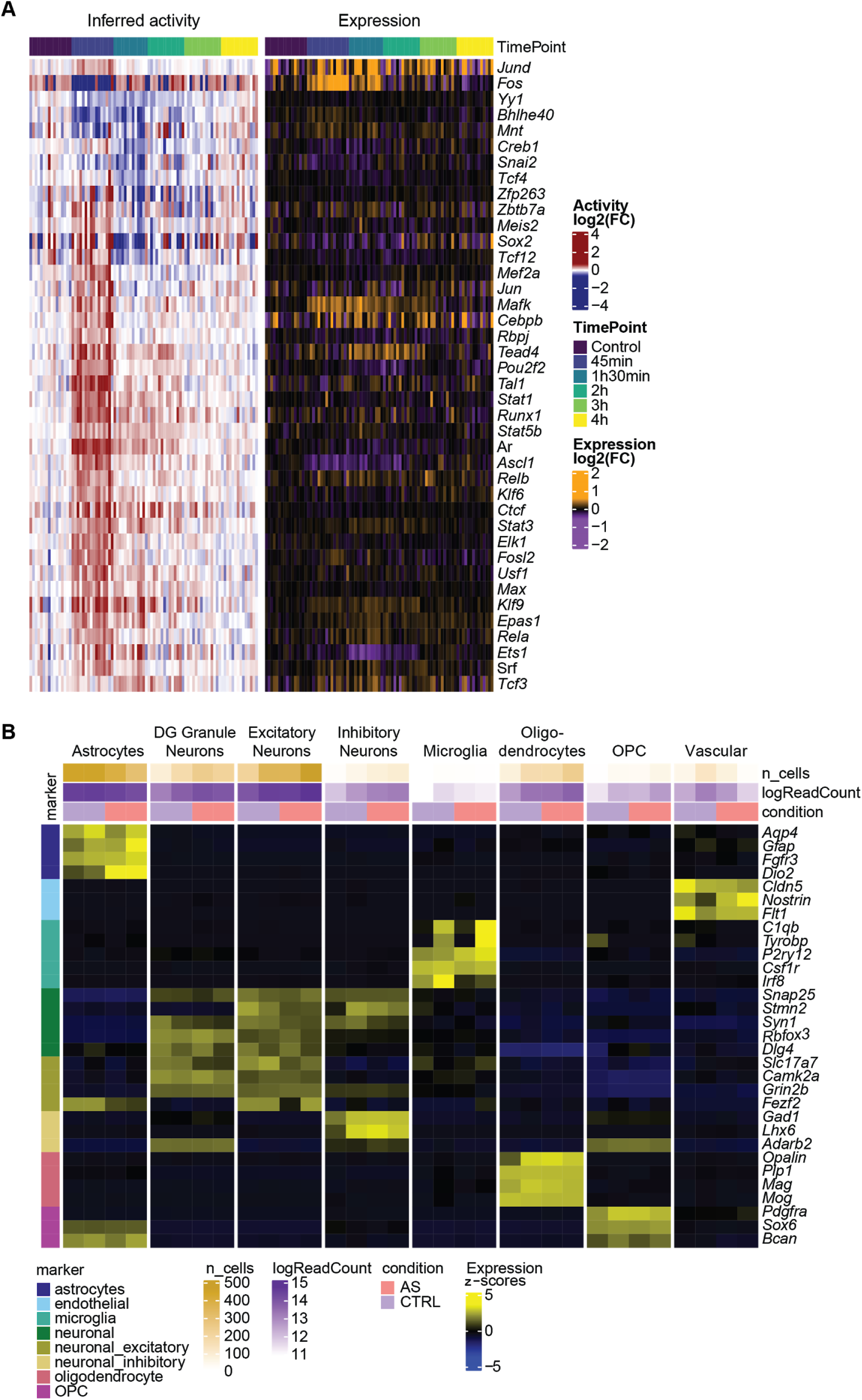
**A)** Relative inferred activity (left) and expression (right) of the transcription factors showing significant changes in activity. **B)** Expression of known markers across the final snRNAseq clusters. Cell-level counts were aggregated by sample and cluster and normalized at the pseudo-bulk level.

**Figure S6:**
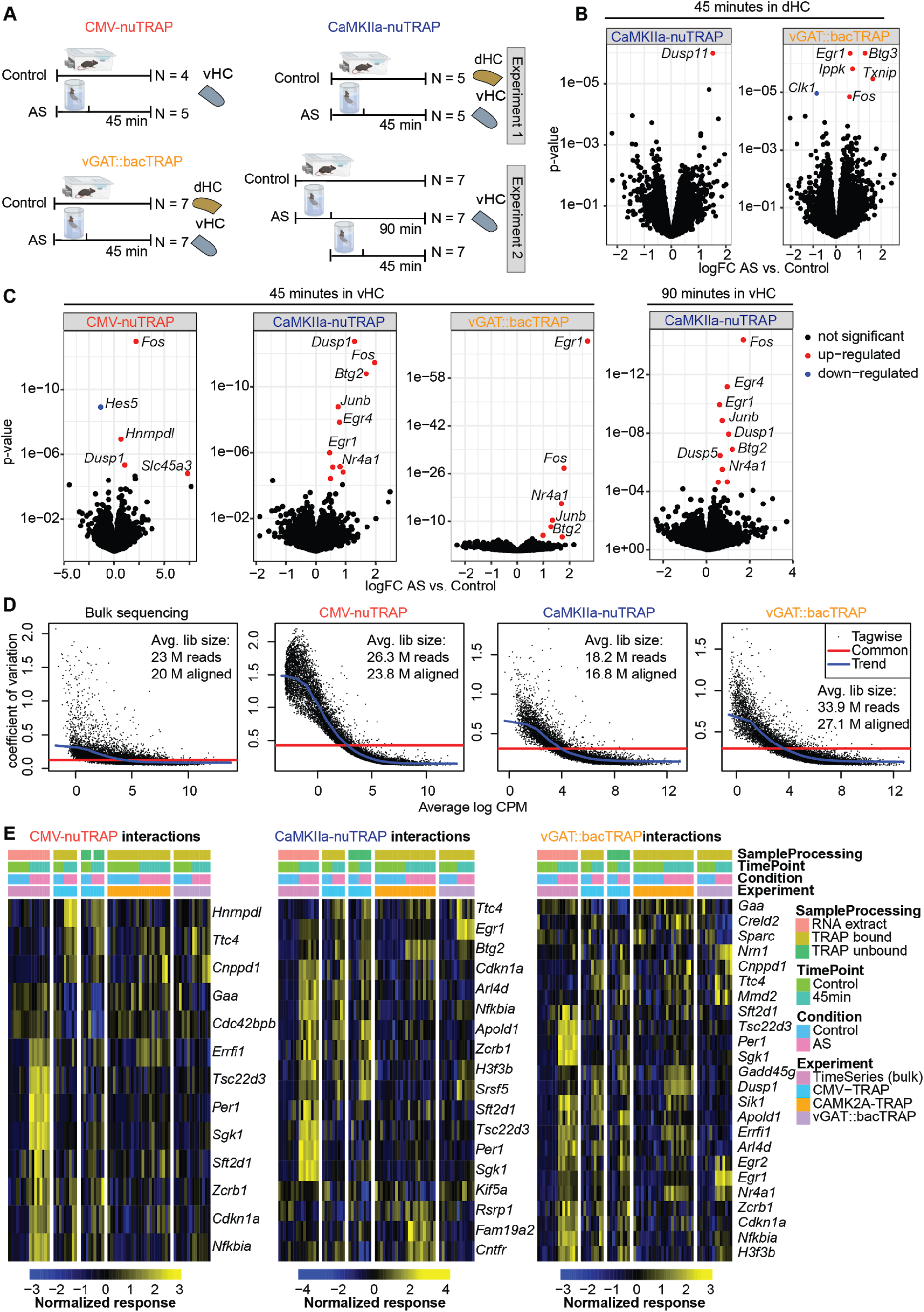
Effects of swim stress on the whole translatome and the translatome of inhibitory and excitatory neurons. **A)** Experimental design of all TRAP experiments. **B)** Volcano plots depicting significant changes in translation in excitatory neurons (CaMKIIa-nuTRAP) and inhibitory neurons (vGAT::bacTRAP) in the dHC at 45 minutes. **C)** Volcano plots depicting significant changes in translation in whole tissue (CMV-nuTRAP), excitatory neurons (CaMKIIa-nuTRAP) and inhibitory neurons (vGAT::bacTRAP) in the vHC at 45 minutes and in excitatory neurons (CaMKIIa-nuTRAP) at 90 minutes. **D)** Plots depicting coefficient of variation (CV) vs logCPM in bulk sequencing, CMV-nuTRAP, CaMKIIa-nuTRAP and vGAT::bacTRAP in the vHC at 45minutes swim vs. controls. **E)** Heatmaps depicting DEGs with significant interaction terms (p interaction <= 0.05) in CMV-nuTRAP, CaMKIIa-nuTRAP and vGAT::bacTRAP and their expression across all datasets

**Figure S7:**
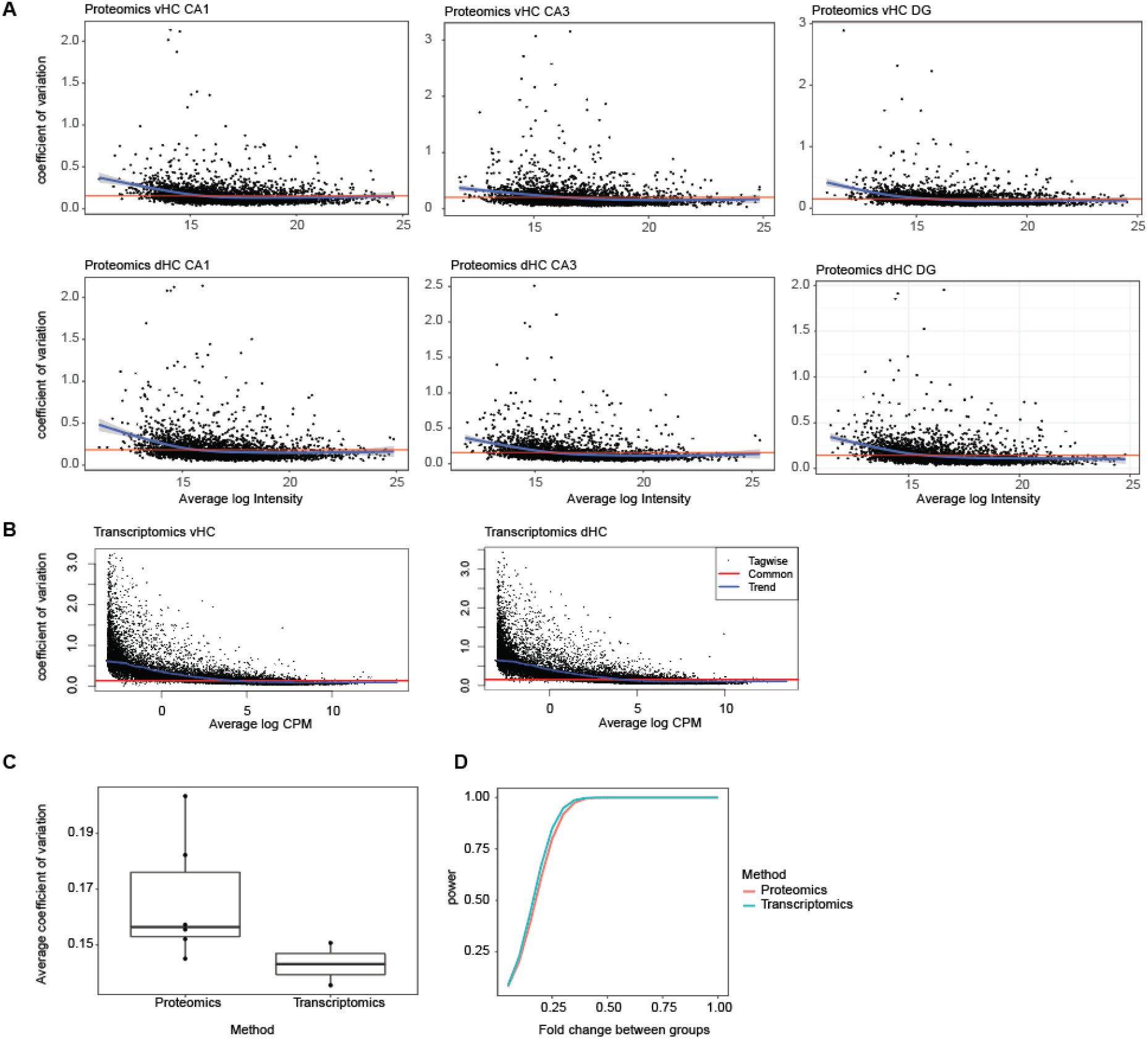
Variability and power of proteomics and transcriptomics analyses. **A)** coefficient of variation (CV) in individual regions and sub-regions using proteomics **B)** CV in individual regions using transcriptomics **C)** Mean CV values from A/B shown as boxplot **D)** Power analysis demonstrating similar sensitivity for both methods in respect to the fold change between groups (calculated with α = 0.05; N = 7 / CV = 0.143 for transcriptomics and N = 8 / CV = 0.166 for proteomics)

